# Poisson representation: a bridge between discrete and continuous models of stochastic gene regulatory networks

**DOI:** 10.1101/2023.07.19.549675

**Authors:** Xinyu Wang, Youming Li, Chen Jia

**Author notes:** These authors contribute equally to this work (Xinyu Wang, Youming Li).

## Abstract

Stochastic gene expression dynamics can be modeled either discretely or continuously. Previous studies have shown that the mRNA or protein number distributions of some simple discrete and continuous gene expression models are related by Gardiner’s Poisson representation. Here we systematically investigate the Poisson representation in complex stochastic gene regulatory networks. We show that when the gene of interest is unregulated, the discrete and continuous descriptions of stochastic gene expression are always related by the Poisson representation, no matter how complex the model is. In addition, using a simple counterexample, we find that the Poisson representation in general fails to link the two descriptions when the gene is regulated. However, for a general stochastic gene regulatory network, we demonstrate that the discrete and continuous models are approximately related by the Poisson representation in the limit of large protein numbers. These theoretical results are further applied to analytically solve many complex gene expression models whose exact distributions are previously unknown.

## 1 Introduction

Over the past two decades, breakthroughs have been made in the theory [1, 2] and experiments [3–5] of single-cell stochastic gene expression dynamics. The models of stochastic gene expression can be classified into two categories: discrete and continuous ones. Discrete models are those where the mRNA and protein numbers change by discrete integer amounts when reactions occur [1, 6]. In continuous models, the mRNA and protein fluctuations correspond to hops on the real axis rather than on the integer axis [7, 8]. Continuous models of stochastic gene expression can be further classified according to whether the fluctuations of genes are modeled discretely or continuously. In previous studies, those models with the gene fluctuations described in the discrete sense and with the mRNA and protein fluctuations described in the continuous sense [9, 10] are sometimes referred to as hybrid models. However, in this study, we will refer to them as continuous models to emphasize the continuous changes of mRNA and protein abundances.

The Poisson representation was first introduced by Gardiner and Chaturvedi [11–13] and serves as a powerful tool to investigate the properties of chemical master equations. Recently, some studies [14–16] used the Poisson representation to examine the relationship between discrete and continuous gene expression models and found that the gene product number distributions of some simple discrete and continuous models are related by the Poisson representation. It has been also shown [17–19] that various continuous models can be viewed as different macroscopic limits of the discrete models.

The importance of the Poisson representation can be seen from the following two specific examples. In experiments, it was observed that many mRNAs and proteins are produced in the bursty sense with the burst size having a geometric distribution [3, 4]. For such bursty gene expression, it has been shown that in steady state, the gene product number of the discrete model has a negative binomial distribution [20], while for the continuous model, the gene product number has a gamma distribution [8]. The discrete negative binomial distribution and the continuous gamma distribution are related by the Poisson representation and thus the former is also referred to as a Poisson-gamma distribution. The second example is the classical telegraph model of stochastic gene expression which describes the synthesis and degradation of the gene product, as well as the switching of the promoter between an active and an inactive state [21]. The discrete telegraph model can be solved exactly [22] and the steady-state gene product number distribution contains a confluent hypergeometric function. The continuous version of the telegraph model was first solved in [23] and the gene product distribution has a beta distribution in steady state. Interestingly, the discrete hypergeometric-type distribution and the continuous beta distribution are also linked by the Poisson representation, and this beautiful relation has been used in the parameter inference of the telegraph model [24].

While the Poisson representation has been shown to be able to link the copy number distributions of some simple discrete and continuous gene expression models [14–16], it remains unclear whether this result can be generalized to complex systems of gene expression and gene regulation. In particular, most all previous studies on the Poisson representation only focus on the expression of a particular gene that is unregulated. The aim of the present paper is to study to whether and what extent the Poisson representation can connect the discrete and continuous models of a general gene regulatory network. Even when the gene of interest is unregulated, it is still not clear whether the Poisson representation works for a general model with complex promoter switching mechanisms, bursty production of mRNAs and proteins, and upstream cellular drives.

The present paper is organized as follows. In Section 2, we recall the definition of univariate Poisson representation and describe the discrete and continuous expression models in detail for an unregulated gene with complex gene state switching, bursty production of protein, degradation of protein, and upstream cellular drives. In Section 3, we show that for an unregulated gene, the protein number distributions for the discrete and continuous models can always be related by the Poisson representation, no matter how complex the model is. In Section 4, we show that the Poisson representation cannot accurately relate the discrete and continuous models for an autoregulated gene; however, it approximately works in the limit of large protein numbers. The results in Sections 3 and 4 are applied to compute the exact or approximate protein number distributions for many complex gene expression models whose analytical solutions are previously unknown. In Section 5, we recall the definition of multivariate Poisson representation and demonstrate that for a general stochastic gene regulatory network, the joint distribution of protein numbers of the discrete and continuous models can be approximately related by the multivariate Poisson representation when protein numbers are large. We conclude in Section 6.

## 2 Poisson representation and models

### 2.1 Poisson representation

We first recall the definition of Poisson representation [11–13]. Let *p*_*n*_, *n* = 0, 1, 2, ⋯ be the probability mass function of a nonnegative integer-valued discrete random variable. We say that *p*_*n*_ has a *Poisson representation* if it can be represented as the integral form

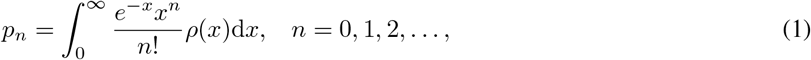

where *ρ*(*x*), *x* ≥ 0 is the probability density function of a nonnegative real-valued random variable. Note that *e*^*−x*^*x*^*n*^*/n*! is the probability mass function of a Poisson distribution with mean *x*. Hence, intuitively, the Poisson representation decomposes a discrete probability distribution *p*_*n*_ into the weights sum of many Poisson distributions with different means *x* ≥ 0, with the weights being distributed according to the probability density *ρ*(*x*). In what follows, we refer to *ρ*(*x*) as the *Poisson kernel* of *p*_*n*_. If Eq. (1) is satisfied, we say that *p*_*n*_ and *ρ*(*x*) are related by the Poisson representation.

Given the discrete distribution *p*_*n*_, it is usually difficult to find its Poisson kernel *ρ*(*x*) directly from Eq. (1). In order to find the Poisson kernel, recall that the generating function *f* (*z*) of the discrete distribution *p*_*n*_ and the Laplace transform 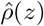 of the Poisson kernel *ρ*(*x*) are defined by

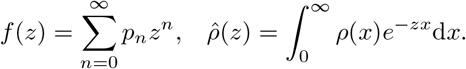

It is well known that any nonnegative integer-valued discrete distribution is uniquely determined by its generating function and any nonnegative real-valued continuous distribution is uniquely determined by its Laplace transform. In terms of the generating function and Laplace transform, Eq. (1) can be converted into the following equivalent form [15]:

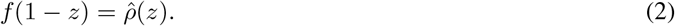

Given the discrete distribution *p*_*n*_, its Poisson kernel *ρ*(*x*) can be determined in the following way. First we compute the generating function *f* (*z*) of *p*_*n*_. If *f* (1 − *z*), as a function of *z*, is the Laplace transform of a nonnegative real-valued random variable, then taking the inverse Laplace transform with respect to *z* gives the Poisson kernel *ρ*(*x*). We stress that while Eqs. (1) and (2) are equivalent, they have different emphases: the former reveals the hidden structure behind the discrete distribution *p*_*n*_, while the latter provides a convenient way to compute the Poisson kernel *ρ*(*x*). It is easy to check that if the Poisson kernel of *p*_*n*_ exists, then *p*_*n*_ must be super-Poissonian, i.e. its variance must be greater than or equal to its mean, and thus its Fano factor must be greater than or equal to 1 [25].

### 2.2 Discrete gene expression models

Here we consider a general stochastic gene expression model with complex gene state switching, bursty production of protein, degradation of protein, and upstream cellular drives (Fig. 1(a)). For simplicity, here we do not take genetic feedback loops into account. The extension to a general gene regulatory network will be discussed in Section 5.

**Figure 1.**
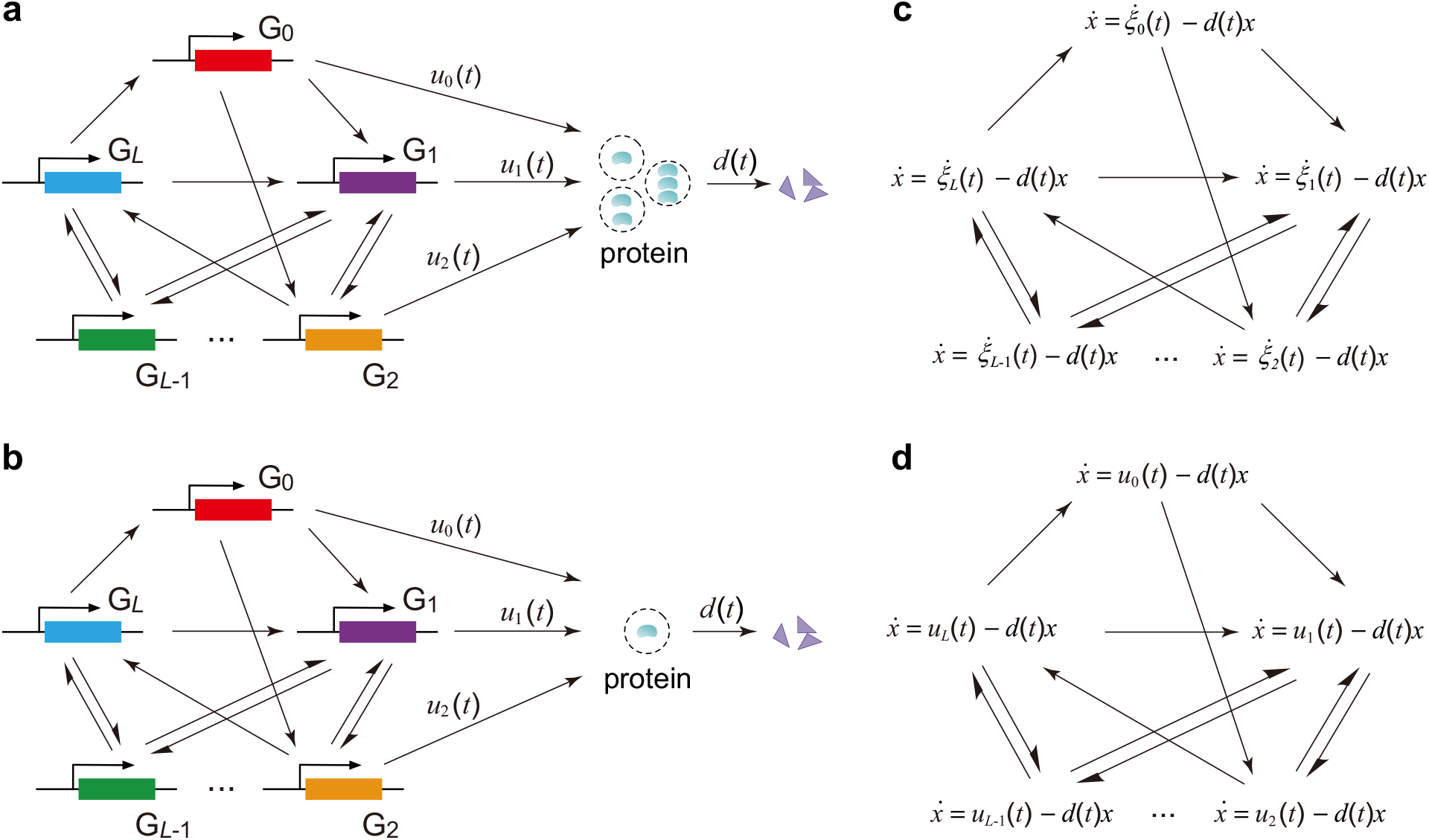
Models. **(a)** Schematic of a gene expression model with complex gene switching mechanism and bursty production of gene product molecules. The gene can exist in *L* + 1 states, *G*_0_, *G*_1_, …, *G*_*L*_, and the switching among them can be arbitrary. Different gene states correspond to different burst frequencies of the gene product (some arrows corresponding to gene product synthesis are omitted in the figure). In each gene state, the gene product molecules are produced in a bursty manner. **(b)** Schematic of a gene expression model with complex gene switching mechanism and non-bursty production of gene product molecules. In each gene state, the gene product molecules are produced in a non-bursty manner. **(c)** Schematic of the switching SDE model, which is a continuous model of stochastic gene expression in the case of bursty production. In each gene state *G*_*i*_, the synthesis and degradation of the gene product is described by the SDE 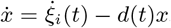. The SDE is driven by a compound Poisson process *ξ*_*i*_(*t*) which describes stochastic gene expression bursts. **(d)** Schematic of the switching ODE model, which is a continuous model of stochastic gene expression in the case of non-bursty production. In each gene state *G*_*i*_, the synthesis and degradation of the gene product is described by the ODE 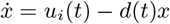.

We assume that the gene of interest can exist in *L* + 1 states, denoted by *G*_0_, *G*_1_, …, *G*_*L*_, and let *P* denote the corresponding protein. These gene states correspond to different conformational states during chromatin remodeling or different binding states with transcription factors [26]. Most previous studies assume that the gene switches between an active and an inactive state with the former having a higher transcription rate than the latter [22]. However, recent experiments suggest that the gene may transition among multiple states with different transcription rates [27–33]. In addition, we assume that the protein is produced in a bursty manner with an arbitrary burst size distribution. Let *n* denote the copy number of protein molecules. Then the gene expression dynamics can be described by the following reactions:

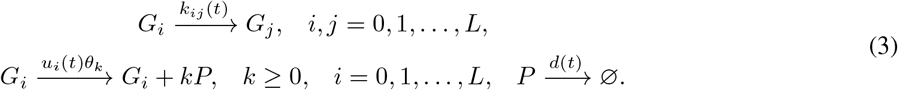

Here we allow all reaction rates to be time-dependent. Such time-varying rates can be understood as upstream cellular drives due to different aspects of cellular environment or gene networks inside the cell [14]. The first reaction describes gene state switching with rates *k*_*ij*_(*t*), the second reaction describes bursty production of protein in all gene states, and the third reaction describes protein degradation with rate *d*(*t*). In particular, in each gene state *G*_*i*_, protein synthesis occurs at a rate *u*_*i*_(*t*) in bursts of a random size sampled from an arbitrary distribution *θ*_*k*_, *k* ≥ 0. This means that in each burst, there is a probability *θ*_*k*_ of producing *k* protein molecules.

The microstate of the gene can be represented by an ordered pair (*i, n*), where *i* = 0, 1, …, *L* is the state of the gene and *n* ≥ 0 is the number of the protein. Let *p*_*i,n*_(*t*) denote the probability of having *n* protein copies when the gene is in state *i*. Then the stochastic gene expression dynamics can be modeled as a Markov jump process, and the evolution of the Markovian model is governed by the following set of chemical master equations (CMEs):

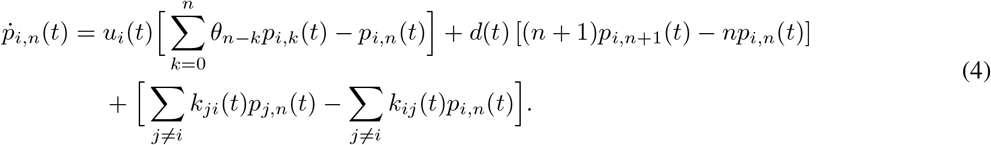

Here the first term on the right-hand side describes protein synthesis, the second term describes protein degradation, and the last term describes gene state switching. Since the gene state and protein number are both discrete variables, the above Markovian model will be referred to as the *discrete model* in what follows.

We emphasize that protein synthesis is not always bursty. Both non-bursty and bursty protein expression are commonly observed in naturally occurring systems [34]. When protein synthesis is bursty, the burst size typically has the geometric distribution *θ*_*k*_ = *p*^*k*^(1 − *p*), which is supported by experiments [4]. The geometric burst size distribution is due to rapid synthesis of protein from a single short-lived transcript [35, 36]. Some other burst size distributions, such as the Poisson and negative binomial distributions, have also been studied in the literature [37–40].

When gene expression is non-bursty, the burst size distribution *θ*_*k*_ = *δ*_1,*k*_ is the point mass at one, where is *δ*_1,*k*_ the Kronecker delta which takes the value of 1 when *k* = 1 and the value of 0 otherwise. In this case, the reactions describing the protein dynamics are given by (Fig. 1(b))

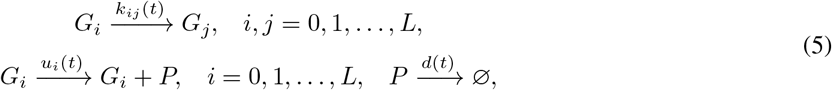

where *u*_*i*_(*t*) is the protein synthesis rate in the gene state *G*_*i*_. The evolution of the discrete model in the non-bursty case is governed by the following set of CMEs:

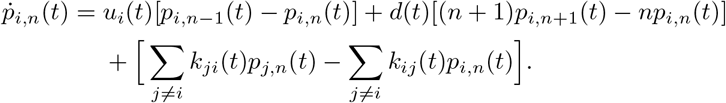

### 2.3 Continuous gene expression models

In many previous papers [8, 18, 40, 41], the gene state is modeled as a discrete variable, while the protein number is modeled as a continuous variable. One important reason for this modeling choice is that in bulk experiments and many single-cell experiments without single-molecule resolution such as flow cytometry, data are often recorded as continuous measurements at a macroscopic scale [18]. The models with discrete gene state and continuous protein abundance are sometimes called hybrid models [42]. However, in the present paper, such kind of models will be referred to as *continuous models* to emphasize that the protein abundance varies continuously.

We have seen that the gene expression dynamics can be modeled discretely as a Markov jump process. We next focus on the continuous counterpart of the discrete model. In the continuous model, the bursty production of protein in each gene state is usually described by a compound Poisson process [40]. This is because in each gene state, transcripts are produced one by one and thus mRNA synthesis can be modeled as a simple Poisson process. Once a transcript is synthesized, it can produce a large number of protein molecules before it is finally degraded. Hence protein synthesis can be modeled as a compound Poisson process. In each gene state *G*_*i*_, the gene expression dynamics can be modeled continuously by the stochastic differential equation (SDE) [18, 19]

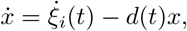

where *x* denotes the protein abundance which varies continuously, the first term on the right-hand side describes bursty protein synthesis with *ξ*_*i*_(*t*) being a compound Poisson process with arrival rate *u*_*i*_(*t*) and jump distribution *w*(*x*), and the second term describes protein degradation with rate *d*(*t*). Here *u*_*i*_(*t*) is the burst frequency and *w*(*x*) is the burst size distribution. Note that in the discrete model, the burst size distribution is *θ*_*k*_, where *k* = 0, 1, 2, … varies discretely, while in the continuous model, the burst size distribution should be chosen differently as *w*(*x*), where *x* ≥ 0 varies continuously. The relationship between *θ*_*k*_ and *w*(*x*) will be explained later.

Since the gene can switch between different states, the gene expression dynamics can be modeled continuously as a hybrid switching SDE [18], as illustrated in Fig. 1(c). Let *p*_*i*_(*x, t*) denote the probability density of the protein abundance *x* when the gene is in state *i*. Then the evolution of the switching SDE model is governed by the Kolmogorov forward equation

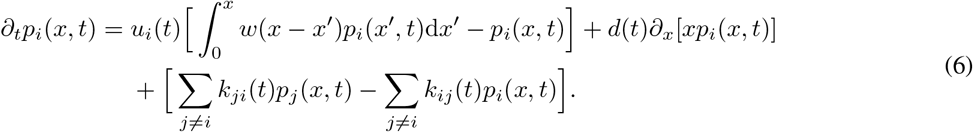

Here the first term on the right-hand side describes protein synthesis, the second term describes protein decay, and the last term describes gene state switching.

The remaining question is how to choose the burst size distribution *w*(*x*) of the continuous model. To answer this, recall that in previous papers, the burst size distribution of the discrete model is often chosen to be the geometric distribution *θ*_*k*_ = *p*^*k*^(1 − *p*) [20], while the burst size distribution of the continuous model is often chosen to be the exponential distribution *w*(*x*) = *e*^*−x/B*^ */B* [8], where 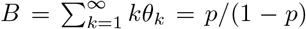 is the mean burst size of the discrete model. A crucial observation is that the geometric and exponential distributions are related by the Poisson representation, i.e.

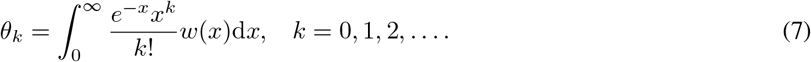

In other words, the exponential distribution is exactly the Poisson kernel of the geometric distribution. Thus in what follows, *we always assume that θ*_*k*_ *and w*(*x*) *are related by the Poisson representation*, even when *θ*_*k*_ is not geometrically distributed.

Note that non-geometric burst size distributions were also studied in some previous papers. In [37], the burst size of the discrete model is assumed to have the Poisson distribution *θ*_*n*_ = *e*^*−λ*^*λ*^*n*^*/n*!. According to our assumption, the corresponding burst size distribution of the continuous model is the point mass at *λ*, i.e. *w*(*x*) = *δ*(*x* − *λ*). In [39], the burst size of the discrete model is assumed to have the negative binomial distribution

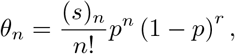

where (*s*)_*n*_ = *s*(*s* + 1) ⋯ (*s* + *n* − 1) is the Pochhammer symbol. In this case, the corresponding burst size distribution of the continuous model is the gamma distribution

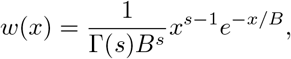

with *B* = *p/*(1 − *p*), since the gamma distribution is the Poisson kernel of the negative binomial distribution.

We next consider the special case where protein synthesis is non-bursty. In this case, in each gene state *G*_*i*_, we use the following ordinary differential equation (ODE) to model the gene expression dynamics [15, 18]:

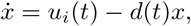

where the two terms on the right-hand side characterize synthesis and degradation of protein. Since the gene can switch between different states, the gene expression dynamics can be modeled continuously as a hybrid switching ODE [43], as depicted in Fig. 1(d). The switching ODE is also called a piecewise deterministic Markov process [44], and this model has been studied extensively in the literature [45–47]. In the non-bursty case, the evolution of the switching ODE model is governed by the Kolmogorov forward equation

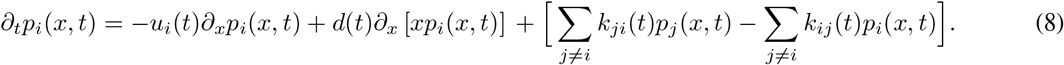

This switching ODE model is also referred to as “effective upstream drive” in [14], since it integrates the effects of upstream cellular drives with time-dependent synthesis and degradation rates.

In fact, Eq. (8) can also be viewed as a special case of Eq. (6). To see this, recall that in the discrete model, the burst size distribution *θ*_*k*_ = *δ*_1,*k*_ is the point mass at one when gene expression is non-bursty. The corresponding burst size distribution for the continuous model should be chosen as

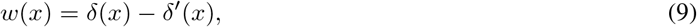

where *δ*(*x*) is Dirac’s delta function and *δ*^*′*^(*x*) is the weak derivative of *δ*(*x*). Under this choice, *θ*_*k*_ and *w*(*x*) are related by the Poisson representation since the generating function of the former is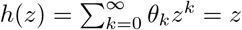, while the Laplace transform of the latter is 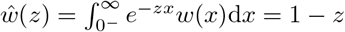 [48], where 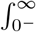 stands for 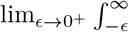. Note that when integrating the delta function, *x* = 0 has to be an interior point of the integration region and hence 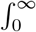 in the definition of the Laplace transform should be modified slightly as 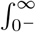. It thus follows from Eq. (2) that *w*(*x*) is the Poisson kernel of *θ*_*k*_. Inserting Eq. (9) into Eq. (6), we immediately obtain Eq. (8).

### 2.4 Examples of Poisson representation

For some simple gene expression systems, it was already known that the steady-state protein number distributions for the discrete and continuous models are related by the Poisson representation [14, 15]. To help readers gain deeper insights, we next provide several examples.

**Example 1**. Let *G* be the gene of interest and let *P* be the corresponding gene product (mRNA or protein). The simplest gene expression model is given by

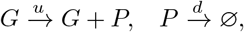

where the gene product is produced in a non-bursty manner. When the system reaches steady state, the gene product number for the discrete Markovian model has the Poisson distribution 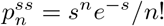, where *s* = *u/d*. On the other hand, since gene expression is non-bursty, the corresponding continuous model is simply the ODE 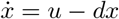, which has a unique globally asymptotic stable equilibrium *s* = *u/d*. In other words, the steady-state distribution of the continuous model is the point mass at *s*, i.e. *p*^*ss*^(*x*) = *δ*(*x* − *s*). It is easy to see that the steady-state solutions for the two models are related by the Poisson representation, i.e.

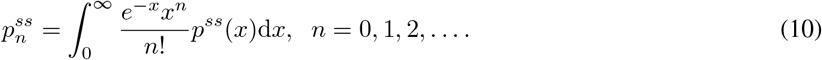

**Example 2**. Consider a gene expression system described by the reactions

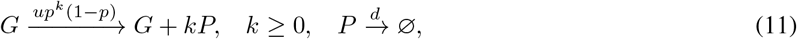

where the gene product is produced in bursts with the burst size having a geometric distribution *θ*_*k*_ = *p*^*k*^(1 − *p*). In steady state, it has been shown [20] that the gene product number for the discrete Markovian model has the negative binomial distribution

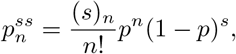

where *s* = *u/d* and (*s*)_*n*_ = *s*(*s* + 1) … (*s* + *n* − 1) is the Pochhammer symbol. Since gene expression is bursty, the corresponding continuous model is the SDE 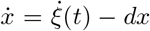, with *ξ*(*t*) being a compound Poisson process with burst frequency *u* and exponential burst size distribution *w*(*x*) = *e*^*−x/B*^*/B*. It is well-known [8] that the gene product number for the continuous SDE model has the gamma distribution

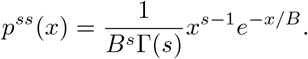

It is easy to check that the steady-state distributions for the two models are related by the Poisson representation.

In single-cell RNA sequencing experiments, the observed number of mRNA reads usually has a negative binomial distribution [49]. The reaction scheme (11) provides a mechanistic understanding of the negative binomial distribution from the biophysical perspective. However, in the biostatistical and bioinformatic literature, the origin of the negative binomial distribution is often interpreted as follows [50–54]: there is a true number *x* of transcripts that varies in the population and is distributed as a gamma random variable. The number of observed reads is distributed as a Poisson random variable with mean *x*. Hence the total number of the reads obeys the Poisson-gamma distribution, i.e. the negative binomial distribution.

**Example 3**. The classical telegraph model of gene expression is described by the following reactions:

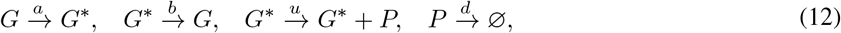

where the gene switches between an inactive state *G* and an active state *G*^*\**^, and the gene product is produced in a non-bursty manner when the gene is active. It has been shown [22] that the steady-state copy number distribution for the discrete model is given by

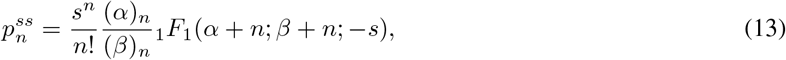

where *α* = *a/d, β* = (*a* + *b*)*/d, s* = *u/d*, and _1_*F*_1_(*α*; *β*; *z*) is the confluent hypergeometric function. Since gene expression is non-bursty, the corresponding continuous model is the switching system of ODEs

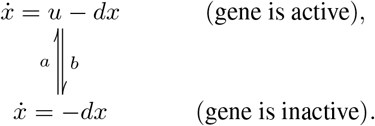

It is also known [9] that the steady-state distribution for the continuous model is the beta distribution

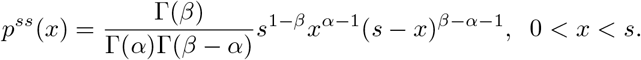

It is easy to check that the generating function *f*^*ss*^(*z*) of the discrete distribution 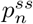 and the Laplace transform 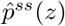 of the continuous distribution *p*^*ss*^(*x*) are related by 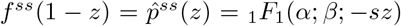. This indicates that the two distributions are related by the Poisson representation. Due to this reason, the steady-state distribution (13) of the telegraph model is also called the Poisson-beta distribution [24, 55–57].

## 3 Poisson representation for unregulated genes

### 3.1 Main results

From the above examples, it can be seen that for some simple gene expression systems, even for systems with a geometrical burst size distribution (Example 2) or two-state gene state switching (Example 3), the steady-state copy number distributions for the discrete and continuous models are related by the Poisson representation. A natural question is whether this result holds for general gene expression systems — does it also hold for systems with an arbitrary burst size distribution, complex multi-state gene switching, and even feedback regulation? In addition, it is also unclear whether the above result holds for the time-dependent dynamic behavior of the system — are the time-dependent distributions for the discrete and continuous models also related by the Poisson representation, even for systems with upstream cellular drive?

In fact, these questions have been partially answered in some previous papers [14–16]. However, the conclusion is not general due to the following reasons. First, all these papers only consider unregulated genes and did not take feedback controls into account. Second, most previous papers [14, 15] only focus on non-bursty gene expression and do not take transcriptional or translational bursting into account; while [16] focuses on bursting, it does not include gene state switching. Third, [15, 16] do not consider upstream cellular drives, while [14] essentially assumes that initially there is no gene product molecules in the cell. In this paper, we will answer the above questions for general gene regulatory networks with complex gene state switching, arbitrary burst size distributions, arbitrary feedback controls, and upstream cellular drives. In addition, we will consider both the steady-state and time-dependent behaviors of the system.

We first focus on the dynamics for unregulated genes, where all the reaction rates are independent of the protein number *n*. The system can be described by the following reactions:

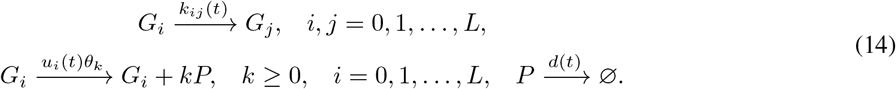

In particular, if *θ*_*k*_ = *δ*_1,*k*_ is the point mass at one, then gene expression in each state is non-bursty. The following theorem is the main result of this section (see Appendix A for the detailed proof).

**Theorem 1**. Consider the gene expression system described by Eq. (14). Suppose that the burst size distributions for the discrete and continuous models are related by the Poisson representation, i.e. Eq. (7) is satisfied. If for each gene state *G*_*i*_, the initial distribution *p*_*i,n*_(0) of the discrete model and the initial distribution *p*_*i*_(*x*, 0) of the continuous model are related by the Poisson representation, i.e.

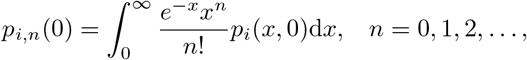

then for each gene state *G*_*i*_ and each time *t* ≥ 0, the time-dependent distributions of the discrete and continuous models are related by the Poisson representation, i.e.

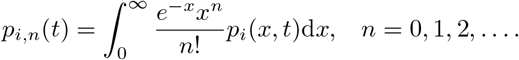

In particular, the steady-state distributions of the discrete and continuous models are also related by the Poisson representation.

The above theorem shows that for unregulated genes, if the burst size distributions and the initial distributions for the discrete and continuous models are related by the Poisson representation, then both the time-dependent and steady-state distributions for the two models are also related by the Poisson representation, no matter how complex the model is.

Theorem 1 has many applications. Over the past two decades, there have been numerous studies focusing on the analytical solutions of various discrete and continuous gene expression models. However, there are still many models whose closed-form solutions are still unknown. Let 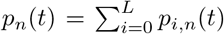 and 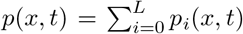 denote the protein number distributions for the discrete and continuous models, respectively. Since the distributions for the two models are related by the Poisson representation, once the exact solution for one model is known, we can use the Poisson representation to recover the exact solution for the other model. Specifically, according to Theorem 1, the generating function *f* (*z, t*) of the discrete distribution *p*_*n*_(*t*) and the Laplace transform 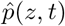 of the continuous distribution *p*(*x, t*) are linked by 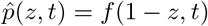. Once the generating function of the discrete model is known, the Laplace transform of the continuous model is also known. Then taking the inverse Laplace transform of *f* (1 − *z, t*) with respect to *z* gives the protein number distribution of the continuous model. On the other hand, once the Laplace transform 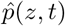 of the continuous model is known, the generating function of the discrete model is given by 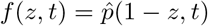. Finally, taking derivatives of *f* (*z, t*) at *z* = 0 gives the protein number distribution of the discrete model, i.e.

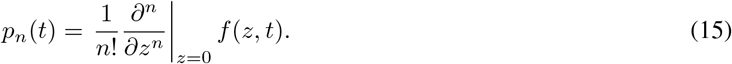

Next we apply the above idea to compute the steady-state and time-dependent distributions for some complex gene expression models. All the exact solutions derived below are novel and have not been obtained in the literature.

### 3.2 Steady-state solution for a multi-state bursty model

Consider a multi-state gene expression system described by the following reactions:

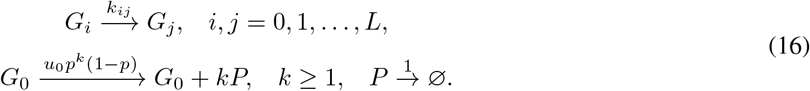

Here we assume that the gene can switch between *L* + 1 different states *G*_0_, *G*_1_, …, *G*_*L*_. The gene has a unique active state *G*_0_ and multiple inactive states *G*_1_, …, *G*_*L*_. The gene product *P* (mRNA or protein) is produced in a bursty manner when the gene is active with the burst size having a geometric distribution, while it is not produced when the gene is inactive. Without loss of generality, we assume that the gene product is degraded with rate *d* = 1. In fact, the discrete model of the above system has been analytically solved in steady-state conditions [58, 59]. However, thus far, the exact steady-state solution for the corresponding continuous model is still unknown.

The consideration behind this model is as follows. Note that in the two-state model (12), the durations of the active and inactive states both have an exponential distribution. This assumption is reasonable for bacterial [60] and yeast [61] cells. However, recent studies have shown that in mammalian cells, the inactive periods for many genes may have a non-exponential distribution with a nonzero peak [27, 28], since the activation of the promoter is a complex multi-step biochemical process due to chromatin remodeling and the binding and release of transcription factors. This indicates that the gene dynamics in the inactive period may contain multiple exponential steps and in sum, the gene would undergo a multiple-state switching process.

Next we compute the steady-state gene product number distribution of the continuous model (see Fig. 2(a) for an illustration), which is a switching SDE. Let *K* = (*k*_*ij*_)_(*L*+1)*×*(*L*+1)_ be the generator matrix of gene state switching, and let *H* be the *L* × *L* matrix obtained by removing the first row and the first column of *K*. We assume that *K* is irreducible, which means that for any pair of gene states *G*_*i*_ and *G*_*j*_, there exists a transition path

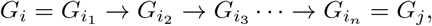

from *G*_*i*_ to *G*_*j*_ with positive switching rates, i.e. 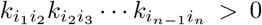. Since *K* is a generator matrix, one eigenvalue of *K* must be zero. Let *α*_1_, …, *α*_*L*_ denote all the nonzero eigenvalues of −*K* and let *β*_1_, …, *β*_*L*_ denote all the eigenvalues of −*H*. It follows from the Perron-Frobenius theorem and the irreducibility of *K* that *α*_*i*_ and *β*_*i*_ all have positive real parts [15]. According to [58, 59], the generating function of the steady-state distribution 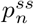 of the discrete model is given by

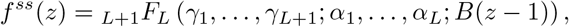

where *B* = *p/*(1 − *p*) is the mean burst size and *γ*_1_, …, *γ*_*L*+1_ are constants satisfying the equations

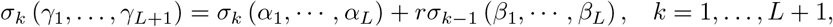

with *σ*_*k*_ being the *k*th elementary symmetric polynomials defined by

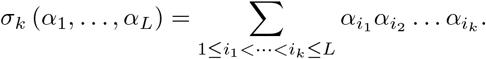

**Figure 2.**
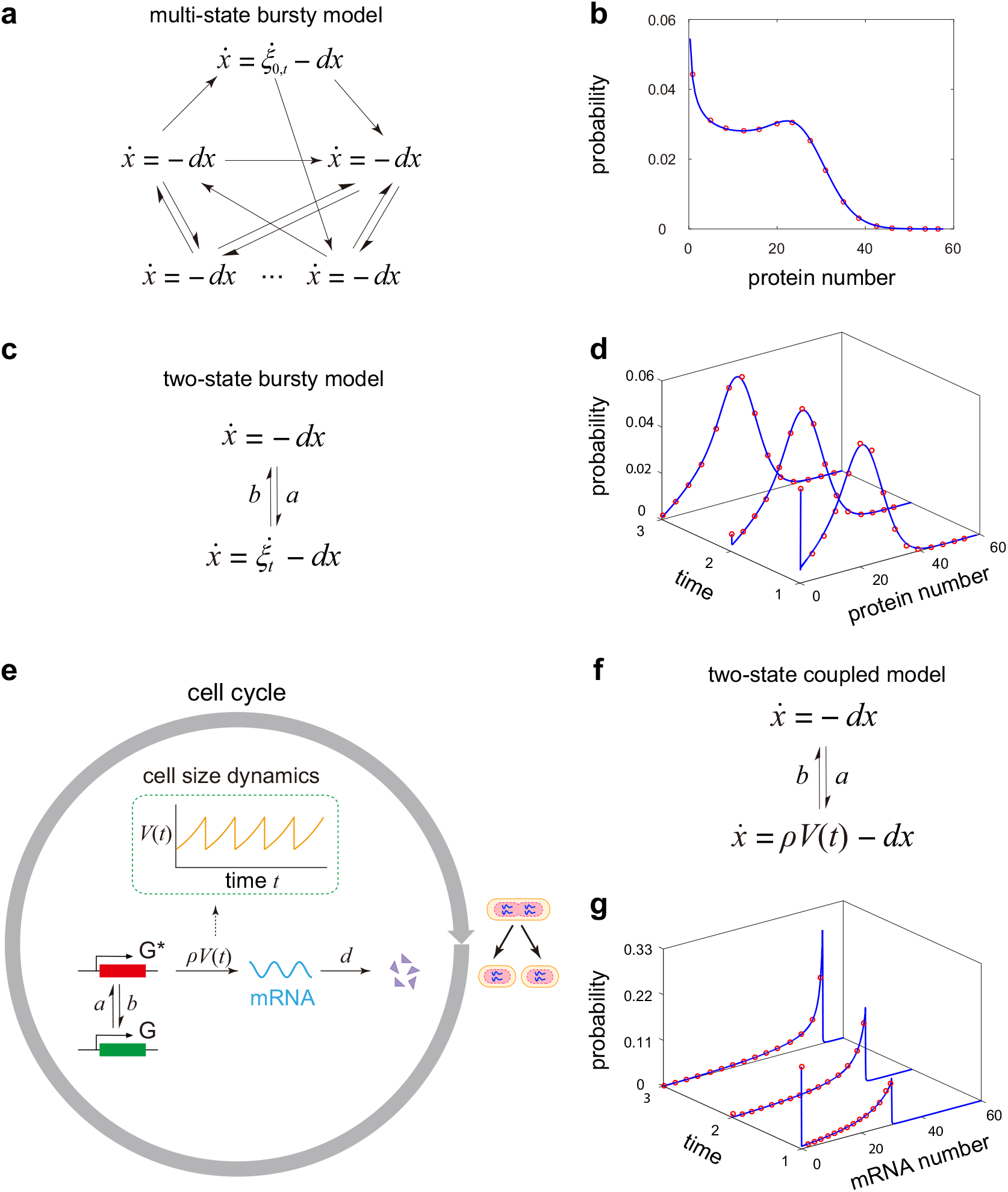
Analytical solutions for three continuous models of stochastic gene expression. **(a)** A multi-state bursty model of gene expression. The switching between multiple states can be arbitrary, and in each gene state, the gene product is produced in a bursty manner with the burst size having an exponential distribution. **(b)** Comparison between the exact steady-state distribution of the continuous model given by Eq. (17) (blue curves) and the numerical one obtained from the SSA (red circles) for the model illustrated in (a). The model parameters are chosen as *L* = 2, *u*_0_ = 30, *B* = 1, *d* = 1, *k*_12_ = 0.35, *k*_13_ = 0.35, *k*_21_ = 0.8, *k*_23_ = 0.35, *k*_31_ = 0.8, *k*_32_ = 0.35. **(c)** A two-state bursty model of gene expression. In the active state, the gene product is produced in a bursty manner with the burst size having an exponential distribution. **(d)** Comparison between the exact time-dependent distributions of the continuous model given by Eq. (18) (blue curves) and the numerical ones obtained from the SSA (red circles) for the model illustrated in (c). The parameters are chosen as *u* = 30, *B* = 1, *d* = 1, *a* = 3.4, *b* = 0.5. The time points are chosen to be *t* = 1, 2, 3. **(e)** Schematic of a coupled dynamic model of gene expression and cell size. Within each cell cycle, the cell size grows exponentially and we use an extended telegraph model to describe gene expression fluctuations with the synthesis rate scaling with cell size. **(f)** The coupled model of gene expression and cell size within each cell cycle. **(g)** Comparison between the exact time-dependent distributions of the continuous model given by Eq. (20) (blue curves) and the numerical ones obtained from the SSA (red circles) for the model illustrated in (e). The parameters are chosen as *d* = 1, *g* = 0.1, *ρ* = 30, *V*_*b*_ = 1, *a* = 2.85, *b* = 0.5, *t* = 1, 2, 3. In (d) and (g), the initial gene product number is zero and the gene initially starts from the inactive state.

From Theorem 1, the steady-state gene product number distribution *p*^*ss*^(*x*) of the continuous model has the Laplace transform 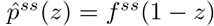. Taking the inverse Laplace transform and using [62, Eq. 3.36.1.1], we obtain

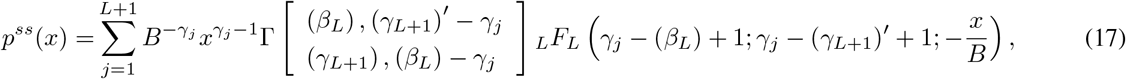

where

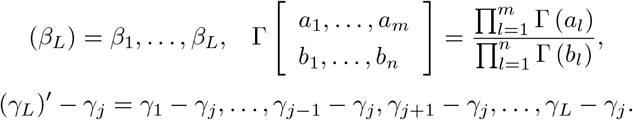

In the special case of *L* = 1, the system reduces to a two-state model and the above result coincides with the one obtained in [18]. To test our analytical solution, we compare it with the numerical solution computed via the stochastic simulation algorithm (SSA) [63] in the case of *L* = 2 under a set of biologically relevant parameters (Fig. 2(b)). Clearly, the copy number distribution is bimodal and the exact solution coincides perfectly with the numerical one.

### 3.3 Time-dependent solution for a two-state bursty model

Consider a two-state gene expression system described by the following reactions:

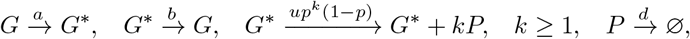

where the gene switches between an active state *G* and an inactive state *G*^*\**^, and the gene product is produced in a bursty manner when the gene is active with the burst size having a geometric distribution. In fact, the discrete model of the above system has been analytically solved in time [59, 64]. However, thus far, the exact time-dependent solution for the corresponding continuous model is still unknown.

We then focus on the time-dependent gene product number distribution of the continuous model (see Fig. 2(c) for an illustration), which is a switching SDE. The generating function of the time-dependent distribution *p*_*n*_(*t*) of the discrete model has been obtained in [64] and is given by

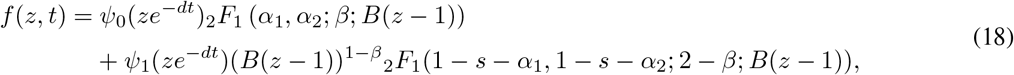

where *B* = *p/*(1 − *p*) is the mean burst size and

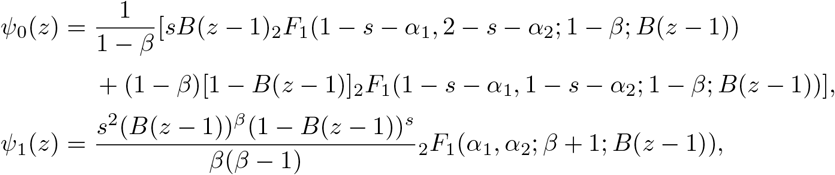

with the parameters *α*_1_, *α*_2_, *β*, and *s* given by

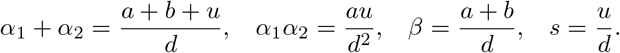

From Theorem 1, the time-dependent gene product number distribution *p*(*x, t*) of the continuous model has the Laplace transform 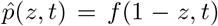. Taking the inverse Laplace transform of *f* (1 − *z, t*) gives the distribution of the continuous model, i.e.

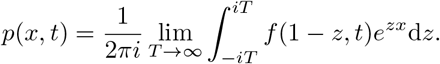

Numerically, we may replace *z* by *iz* in the Laplace transform to obtain the Fourier transform of *p*(*x, t*) and then take the inverse Fourier transform to recover *p*(*x, t*). Fig. 2(d) compares this exact solution with the SSA. Clearly, they are in perfect agreement with each other.

### 3.4 Time-dependent solution for a coupled dynamic model of gene expression and cell size

Thus far, all the analytically solutions obtained are based on models without upstream cellular drive, i.e. models whose reaction rates are time-independent. Next we consider a complex gene expression model with time-dependent synthesis rate. The model couples gene expression dynamics to cell size dynamics and cell cycle events [65–68], and is described as follows (see Fig. 2(e) for an illustration).

1) Let *T* denote the cell cycle duration and let *V* (*t*) denote the cell volume at time *t*. We assume that cell volume grows exponentially within each cell cycle, i.e. 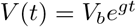 for any 0 ≤ *t* ≤ *T*, where *V*_*b*_ is the cell volume at birth and *g* is the growth rate. The exponential growth of cell volume is commonly observed in various cell types [69]. For simplicity, we assume that the doubling time *T* and the growth rate *g* do not involve any stochasticity.

2) In each cell cycle, we use the following two-state model to describe gene expression dynamics:

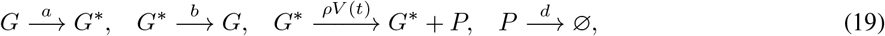

where the gene switches between an active state *G* and an inactive state *G*^*\**^, and the gene product is produced when the gene is active with the synthesis rate scaling with cell volume. In fission yeast, mammalian cells, and plant cells, there is evidence that the mean mRNA or protein number for many genes scales with cell size in order to maintain approximately constant concentrations, a phenomenon known as concentration homeostasis [69, 70]. This is due to a coordination of the synthesis rate with cell volume, as assumed in Eq. (19).

3) At cell division, the mother cell is divided into two daughter cells. The volumes of the two daughter cells are assumed to be the same and exactly one half of the volume before division (of course there is some stochasticity in the partitioning of cell size [71, 72] which we are here ignoring). Moreover, we assume that each gene product molecule has probability 1*/*2 of being allocated to each daughter cell.

Recently, the discrete model of the above system has been solved analytically and the full time-dependent distributions of the gene product number across cell cycles have been obtained [68]. However, the time-dependent solution for the corresponding continuous model is still unknown. For simplicity, here we only focus on the gene expression dynamics within a cell cycle under arbitrary initial conditions. Note that the reaction scheme (19) is a special case of our general model (14) with the synthesis rate being time-dependent. According to [68], the generating function of the gene product number distribution of the discrete model within a cell cycle is given by

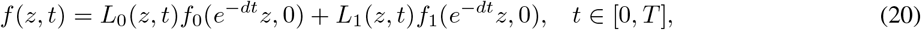

where 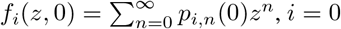 are the generating functions at *t* = 0 which can be determined by the initial conditions, and the functions *L*_*i*_, *i* = 0, 1 are given by

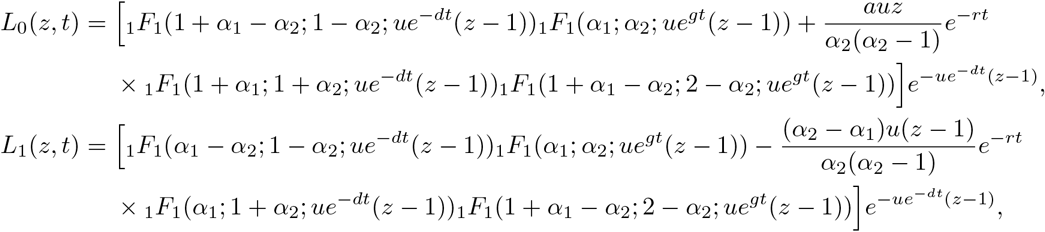

with the parameters *r, α*_1_, *α*_2_, and *u* given by

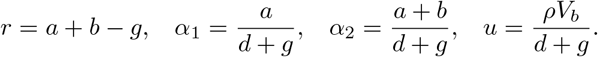

Note that when the growth rate *g* = 0, the synthesis rate *ρV* (*t*) = *ρV*_*b*_ is a constant and the time-dependent solution given in Eq. (20) coincides with that of the classical telegraph model [23, 73].

We then focus on the time-dependent gene product number distribution of the continuous model (see Fig. 2(f) for an illustration), which is a switching SDE. From Theorem 1, the time-dependent distribution *p*(*x, t*) of the continuous model within a cell cycle has the Laplace transform 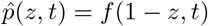. Fig. 2(g) shows that the analytical solution agrees perfectly with the SSA. Here we assume that initially there are no gene product molecules in the cell and the gene is inactive. This mimics the situation where the gene has been silenced by some repressor over a period of time such that all gene product molecules have been removed via degradation; at time *t* = 0, the repressor is removed and we study how gene expression recovers.

## 4 Poisson representation for autoregulated genes

### 4.1 A counterexample

We have seen that for unregulated genes, the gene product number distributions of the discrete and continuous models are always related by the Poisson representation. A natural question is whether this also holds for regulated genes. To answer this, we focus on a multi-state model of autoregulatory gene expression, which is described by the following reactions:

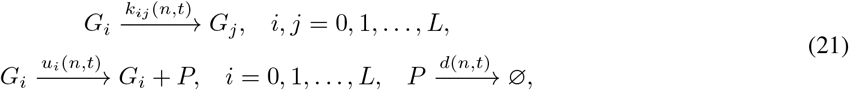

where *G*_0_, *G*_1_, …, *G*_*L*_ are *L* + 1 different gene states and *P* is the corresponding protein. Note that these reactions are the same as those in Eq. (5). Due to autoregulation, following [74], we allow all rate constants to depend on the protein number *n*. If the gene is unregulated, then all rate constants are independent of *n*. For simplicity, here we do not take protein bursting into account. However, the results can be generalized easily to the case of bursty protein synthesis [75].

Next we consider the discrete and continuous models for the autoregulatory system. Let *p*_*i,n*_(*t*) denote the probability of having *n* protein copies when the gene is in state *i*. The evolution of the discrete model is governed by the following set of CMEs:

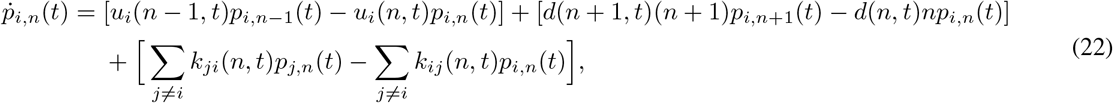

where the three terms on the right-hand side represent protein synthesis, protein degradation, and gene state switching, respectively. For the continuous model, in each gene state *G*_*i*_, we use the following ODE to model the gene expression dynamics:

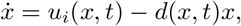

where the two terms on the right-hand side characterize synthesis and degradation of the protein. Since the gene can switch among different states, the gene expression dynamics can be modeled continuously as a hybrid switching system of ODEs. Let *p*_*i*_(*x, t*) denote the probability density of the protein abundance *x* when the gene is in state *i*. The evolution of the switching ODE model is governed by the Kolmogorov forward equation

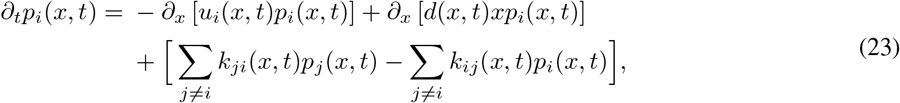

where the three terms on the right-hand side describe protein synthesis, protein degradation, and gene state switching, respectively.

Unfortunately, for regulated genes, the protein number distributions for the discrete and continuous models in general fail to be related by the Poisson representation. This can be seen from the following example.

**Example 4**. Consider a negative autoregulatory gene circuit (see Fig. 3(a) for an illustration) with the following reactions [76, 77]:

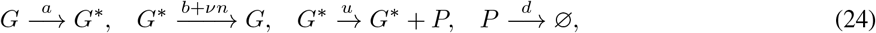

where *G* and *G*^*\**^ denote the inactive and active states of the gene and *P* denotes the corresponding protein. Due to negative autoregulation, the switching rate from *G*^*\**^ to *G* depends on the protein number *n*. Here *a* and *b* are the spontaneous switching rates between the two gene states and *ν* characterizes the strength of negative feedback. It follows from [76, 77] that the steady-state generating function of the protein number distribution for the discrete model is given by

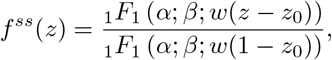

where

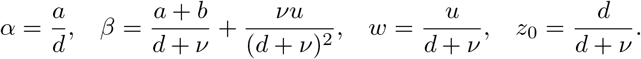

**Figure 3.**
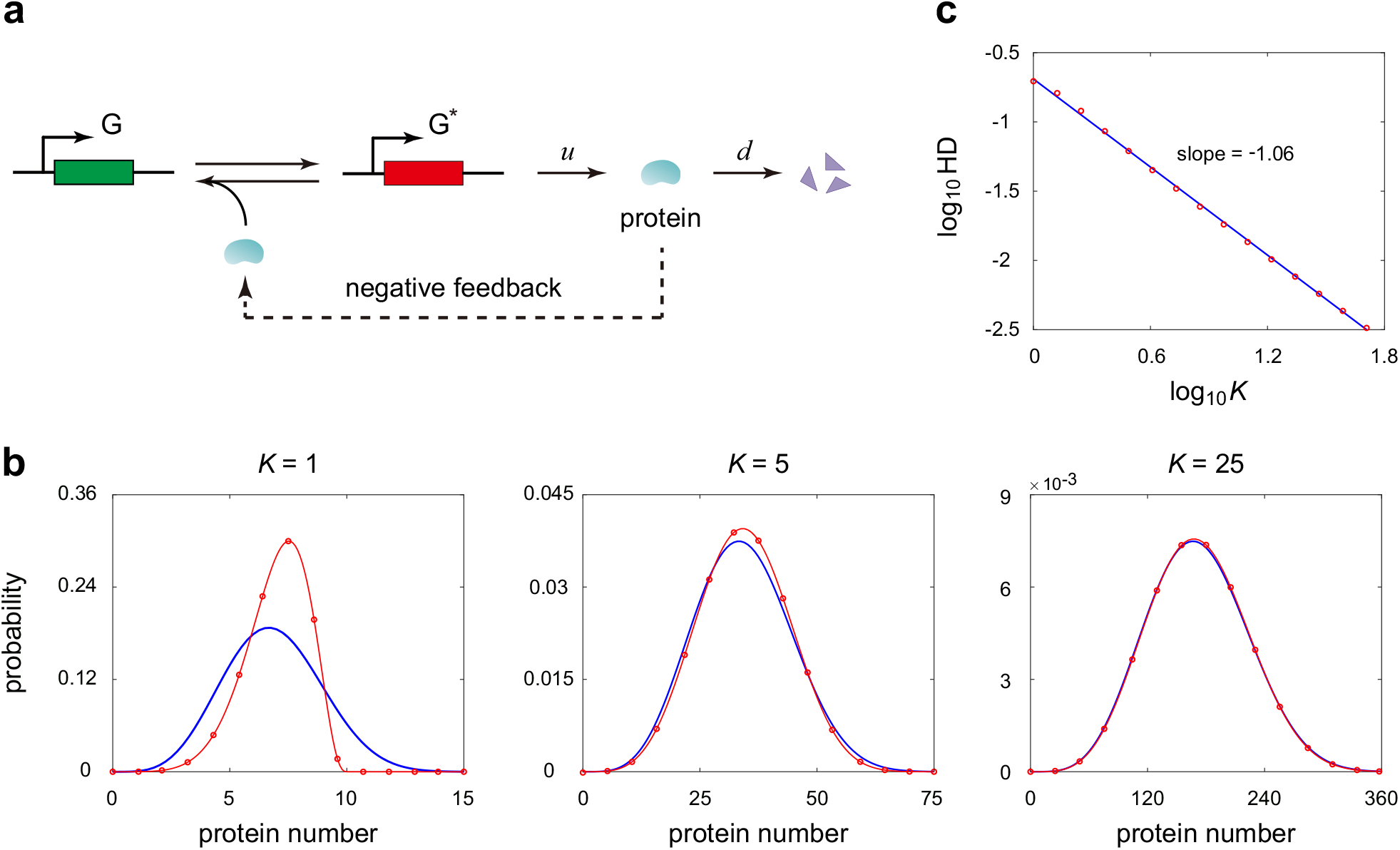
Poisson representation for a negative feedback loop. **(a)** A negative autoregulatory gene circuit, where the binding of the protein to the gene inhibits the expression of itself. **(b)** Comparison of the Poisson kernel *ρ*^*ss*^(*x*) of the discrete model given by Eq. (25) (red circles and red curves) and the exact steady-state distribution *p*^*ss*^(*x*) of the continuous model given by Eq. (26) (blue curves) as the system size *K* increases. The parameters are chosen as *a* = 5, *b* = 2, *d* = 1, *ū* = 25, 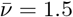. **(c)** The logarithm of the Hellinger distance (HD) between *ρ*^*ss*^(*x*) and *p*^*ss*^(*x*) is approximately a linear function of the logarithm of *K* with slope *−*1.

Since the generating function and the Poisson kernel are related by 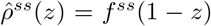 (see Eq. (2)), it is easy to check that the Poisson kernel of the protein number distribution is given by [62, Eq. 3.33.1.1]

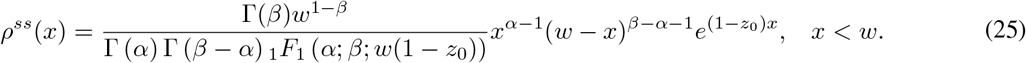

On the other hand, the steady-state protein number distribution of the corresponding continuous model has been computed in [18] and is given by

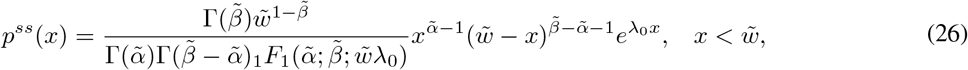

where

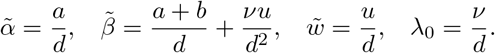

Eqs. (25) and (26) look similar. Comparing these two equations in detail, we find that *ρ*^*ss*^(*x*) ≠ *p*^*ss*^(*x*), which means the steady-state solutions for the discrete and continuous models are not related by the Poisson representation. However, when the feedback strength is much smaller than the degradation rate, i.e. *ν* « *d*, we have

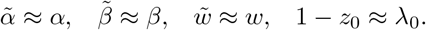

In this case, the steady-state solutions for the two models are approximately related by the Poisson representation. We emphasize that the condition *ν* « *d* is not very strong and is satisfied for a wide range of biological systems. To see this, note that the feedback contribution *νn* to the gene switching rate from *G*^*\**^ to *G* usually has the same order as the spontaneous contribution *b*. This suggests that *b/ν* should have the same order as the mean protein number in the active gene state, i.e. ⟨*n*⟩_active_ = *u/d*. Hence *d/ν* should have the same order as *u/b*. Recent single-cell experiments on transcriptional bursting of mammalian cells have shown that *u/b* is relatively large. In [27], the authors monitored protein expression in mouse fibroblasts using live-cell imaging and found that the values of *a, b*, and *u* for different genes are typically on the order of 0.01/min, 0.1/min, and 1/min, respectively (see Figs. 1(D), 1(E), and S8 of [27] for details). These experimental results imply that the condition *ν* ≪ *d* is indeed satisfied in many biological systems.

From the above example, we have seen that when the gene is regulated, the solutions for the discrete and continuous models are not generally related by the Poisson representation. However, the Poisson representation can still build a bridge between the two models in some special cases. Next we will show that in the limit of large protein numbers, the distributions for the discrete and continuous models are approximately related by the Poisson representation.

### 4.2 Main results

To discuss the limit case of large protein numbers, we introduce a new parameter *K* which stands for the system size, with *K* → ∞ corresponding to a macroscopic scale. In previous papers, the system size *K* is often understood to be the cell volume [5] or the typical protein number in the active gene state [78, 79]. With this new parameter, the limit of large protein numbers corresponds to the limit of *K* → ∞. Since the protein number is mainly controlled by the synthesis rate, it is natural to assume that the protein synthesis rate is proportional to *K*. Specifically, we assume that the synthesis rates in all gene states are proportional to *K*, while the other rate parameters are all functions of the protein concentration, i.e.

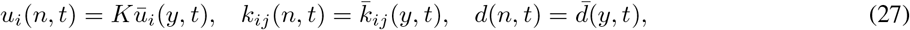

where *y* = *n/K* is the protein concentration, and 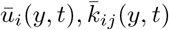, and 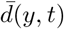 are functions independent of *K*. We will next prove that the protein number distributions for the discrete and continuous models are related by the Poisson representation when *K* ≫ 1.

To this end, we first focus on the dynamics of the protein concentration. Let 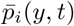 denote the probability density of the protein concentration *y* = *n/K* when the gene is in state *i*. We next derive the dynamic equation of the protein concentration using the discrete model. When *K* ≫ 1, the probability density 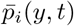 of the protein concentration and the probability distribution *p*_*i,n*_(*t*) of the protein number should be related by [18]

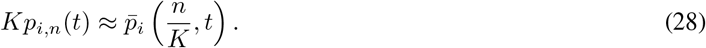

Multiplying *K* on both sides of Eq. (22) and applying Eq. (27), we obtain

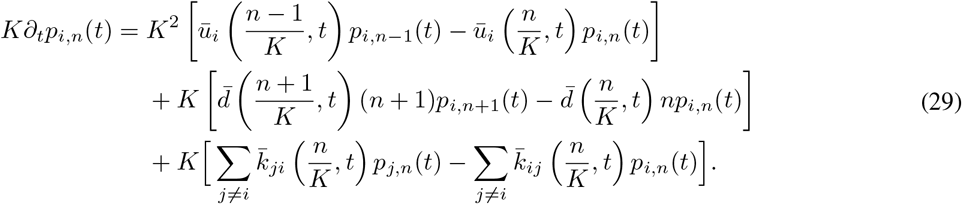

Taking the limit of *K* → ∞ in the above equation and applying Eq. (28), we find that the evolution of the protein concentration is governed by

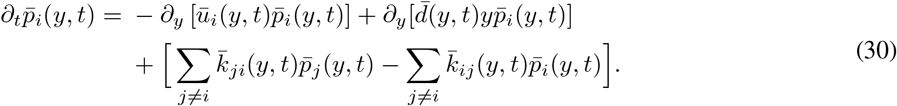

Since *ū*_*i*_(*y, t*) 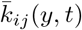, and 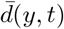 are all independent of *K*, the protein concentration distribution 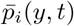 is also independent of *K* when *K* ≫ 1.

Recall that for the continuous model, the evolution of the protein number is described by Eq. (23). Interestingly, comparing Eq. (30) with Eq. (23), we find that the probability density 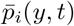 of the protein concentration and the probability density *p*_*i*_(*x, t*) of the protein number are related by

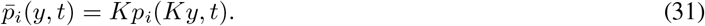

To proceed, let 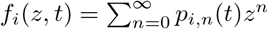 be the generating function of the discrete distribution *p*_*i,n*_(*t*). When *K* ≫ 1, applying the two scaling relations given in Eqs. (28) and (31), we obtain

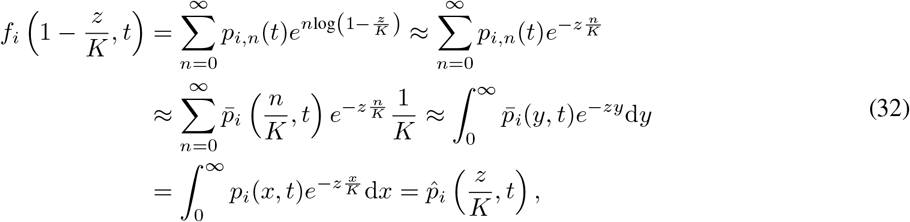

where we have used the fact that log(1 + *x*) ≈ *x* when |*x*| ≪ 1 and the fact that a Riemann sum converges to a Riemann integral as the partition size tends to zero. This shows that the generating function of the discrete distribution *p*_*i,n*_(*t*) and the Laplace transform of the continuous distribution *p*_*i*_(*x, t*) are approximately equal when *K* ≫ 1. Hence, for autoregulatory systems, the time-dependent solutions of the discrete and continuous models are approximately related by the Poisson representation in the limit of large protein numbers, i.e.

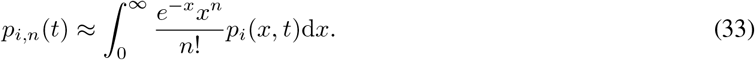

### 4.3 Numerical validation

We have shown that in the limit of large protein numbers, the solutions for the discrete and continuous models for an autoregulatory system are approximately related by the Poisson representation. In other words, the Poisson kernel of the protein number distribution for the discrete model should be close to the distribution for the continuous model. We emphasize that this result is not true when the mean protein number is small. To test our results, we revisit the negative autoregulatory gene circuit described by Eq. (24). Here we assume that the protein synthesis rate *u* is proportional to the system size *K*, i.e. *u* = *Kū*. Since the other rate parameters are all functions of the protein concentration, the feedback strength *ν* should scale with *K* as 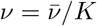. In the limit of large protein numbers, i.e. *K* ≫ 1, it is easy to verify that the parameters given in Eqs. (25) and (26) satisfy

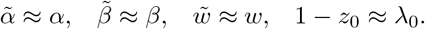

This shows that the Poisson kernel of the protein number distribution for the discrete model is indeed close to the distribution for the continuous model.

Fig. 3(b) compares the Poisson kernel of the discrete distribution given in Eq. (25) and the continuous distribution given in Eq. (26) as *K* increases. Clearly, the two distributions differ significantly when *K* is small, while they are in perfect agreement when *K* is large, as expected. In addition, Fig. 3(c) illustrates the Hellinger distance *H* between the two distributions as a function of *K*. Interestingly, we find that log_10_ *H* is approximately a linear function of log_10_ *K* with slope *κ* ≈ −1, which implies that the Hellinger distance decays at the speed of 1*/K*. This again demonstrates that the Poisson representation becomes valid when protein numbers are large.

### 4.4 Two-state model with complex feedback regulation

We then apply our results to a complex autoregulatory gene expression model whose analytical solution has not been obtained previously. According to our results, if the exact solution of the continuous model *p*(*x, t*) is known, then we can use Poisson representation with Poisson kernel *p*(*x, t*) to construct an approximate solution of the corresponding discrete model when protein numbers are large. Similarly, if the exact solution of the discrete model *p*_*n*_(*t*) is known, then the Poisson kernel of *p*_*n*_(*t*) serves as an approximate solution of the corresponding continuous model in the limit of large protein numbers.

Here we consider the following autoregulatory gene circuit with general feedback regulation:

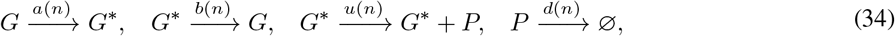

where the protein is produced in a non-bursty manner and all rate constants depend on the protein number *n*. The protein number distribution for this system has not been solved analytically before. Next we will first derive the exact steady-state distribution of the continuous model by generalizing the result in [47], and then obtain the approximate steady-state distribution of the discrete model by using the Poisson representation.

We first obtain the analytical solution of the continuous model. The autoregulatory gene circuit can be modeled continuously as the following switching ODE:

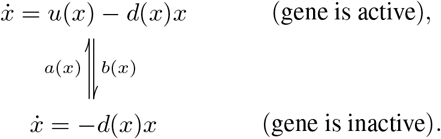

For simplicity, we assume that the ODE 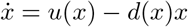 has an unique globally asymptotic stable equilibrium *x*^*\**^with *x* ∈ [0, ∞]. Let *p*_0_(*x, t*) and *p*_1_(*x, t*) denote the probability densities of the protein number when the gene is in the inactive and active states, respectively. Then the evolution of *p*_0_(*x, t*) and *p*_1_(*x, t*) is governed by

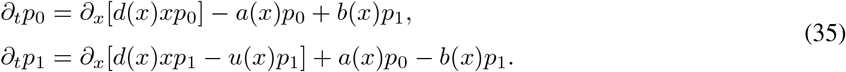

When the system reaches the steady state, summing the two equations in Eq. (35) yields

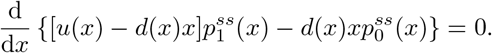

This shows that

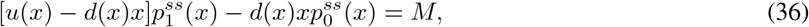

where *M* is a constant. Since 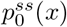 and 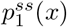 vanish at infinity, we must have *M* = 0. Hence we can define a function

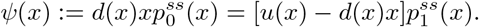

It then follows from Eq. (35) that

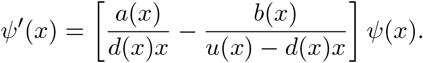

Integrating the above equation yields

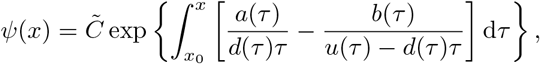

where 0 *< x*_0_ *< x*^*\**^ is arbitrarily chosen and 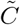 is a normalization constant. Then the steady-state protein number distribution of the continuous model can be solved analytically as

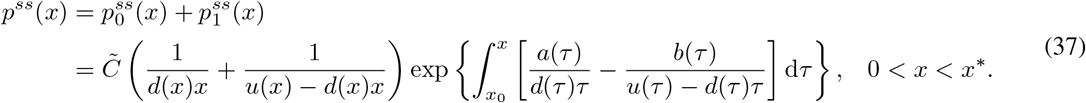

According to our results, the Poisson representation links the discrete and continuous models in the limit of large protein numbers. Hence when protein numbers are large, the steady-state protein number distribution for the discrete model has the following generating function:

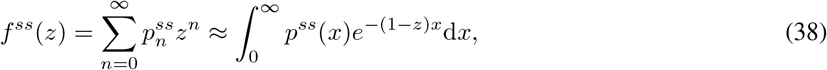

where *p*^*ss*^(*x*) is given by Eq. (37). Taking the derivatives of *f*^*ss*^(*z*) at *z* = 0 gives the approximate steady-state distribution 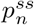 of the discrete model.

To test the accuracy of this solution, we consider a positive autoregulatory feedback loop with the switching rates having the functional forms of *a*(*n*) = *a*_0_ + *a*_1_*n*^2^ and *b*(*n*) = *b*. This corresponds to the case where the switching from the inactive to the active state requires the cooperative binding of two protein molecules (Fig. 4(a)). In addition, we assume that the synthesis rate *u*(*n*) = *u* is independent of *n*, and the protein decays via the Michaelis-Menten propensity function 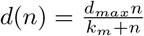 with *d*_*max*_ being the maximum degradation rate [64]. This describes enzyme-catalyzed protein degradation. In this case, the steady-state distribution of the continuous model is given by (see Appendix B for details)

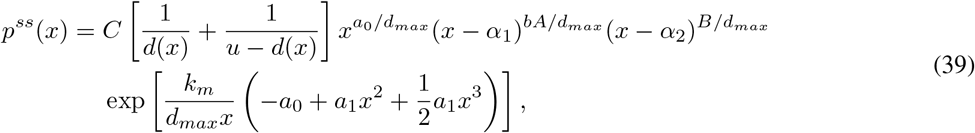

where *C* is a normalization constant and the parameters *α*_1_, *α*_2_, *A*, and *B* are given by

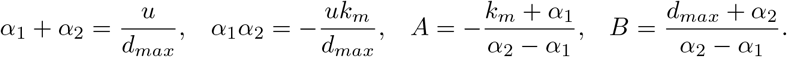

**Figure 4.**
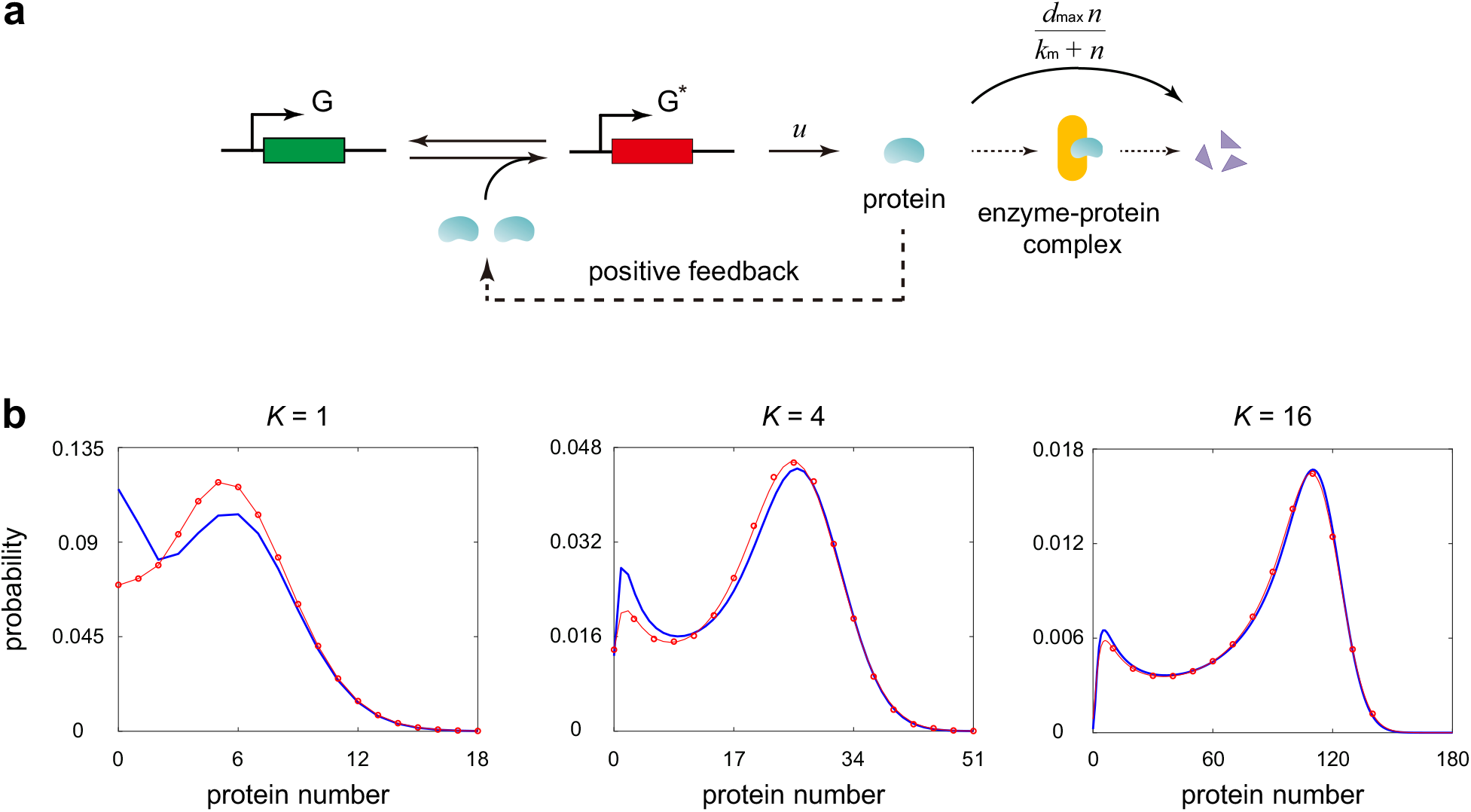
Poisson representation for a positive autoregulatory gene circuit with cooperative binding and enzyme-catalyzed degradation. **(a)** A positive autoregulatory gene circuit. Two protein molecules cooperatively bind to the gene to activate its own expression. The degradation of the protein is enzyme-catalyzed with the rate having the Michaelis-Menten form. **(b)** Comparison of the approximate theoretical distribution of the discrete model given by Eqs. (38) and (39) (red circles and red curves) and the numerical one computed computed using FSP (blue curves). When computing the theoretical distribution of the discrete model, we used the discrete Fourier transform algorithm proposed in [81]. This algorithm can recover the distribution directly from the generating function without calculating the higher order derivatives as in Eq. (15), which is very time-consuming and numerically unstable. The parameters are chosen as *a*_0_ = 1, *ā*_1_ = 0.2, *b* = 2, *ū* = 15, *d*_*max*_ = 2.25, 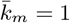.

Finally, we introduce the system size *K* and assume that the parameters *u, a*_1_, and *k*_*m*_ scale with *K* as

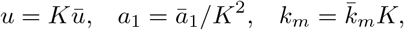

where *ū, ā*_1_, and 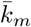 are constants independent of *K*. Then Eq. (38) gives the approximate steady-state distribution of the discrete model when *K* ≫ 1. Fig. 4(b) compares this approximate solution with the numerical one obtained using the finite-state projection algorithm (FSP) [80]. When performing FSP, we truncate the state space at a large integer *N* and solve the truncated master equation numerically. The truncation size is chosen as *N* = 3*Kū/d*, which guarantees that the probability that the protein number is outside the truncated size is very small and practically can always be ignored. It is clear that the approximate solution deviates from the numerical one for small *K*, but it becomes increasingly accurate as *K* becomes larger.

A phenomenon that deserves special attention is that under enzyme-catalyzed protein degradation, the system is capable of producing a bimodal distribution with two non-zero peaks (middle and right panels in Fig. 4(b)). Note that here protein molecules cannot be produced when the gene is in the inactive state. If the protein is degraded via first-order kinetics with constant rate, then previous studies have shown that when bimodality occurs, the bimodal protein number distribution must have both a zero mode and a non-zero mode, and it can never produce two non-zero modes, even when cooperative protein binding to the gene is taken into account [82]. This suggests that the two non-zero peaks in the protein number distribution observed in Fig. 4(b) is due to Michaelis-Menten-type degradation, rather than due to cooperative protein binding.

## 5 Generalization to general gene regulatory networks

### 5.1 Multivariate Poisson representation

In previous sections, we have discussed the Poisson representation for systems with only one gene. Here we will extend the results to a general gene regulatory network, which may involve multiple genes. To this end, we recall the definition of multivariate Poisson representation.

Let 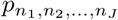 be the joint probability distribution of a nonnegative integer-valued discrete random vector. We say that 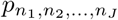 has a Poisson representation if it can be represented in the following form:

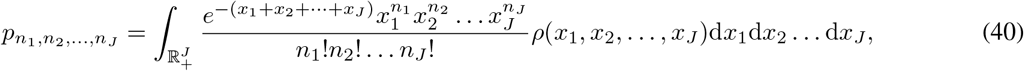

where *ρ*(*x*_1_, *x*_2_, …, *x*_*J*_) is the joint probability density of a nonnegative continuous random vector. In what follows, we refer to *ρ*(*x*_1_, *x*_2_, …, *x*_*J*_) as the Poisson kernel of 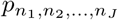. If Eq. (40) is satisfied, we also say that 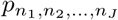 and *ρ*(*x*_1_, *x*_2_, …, *x*_*J*_) are related by the Poisson representation.

The generating function of the discrete distribution 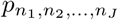 is defined by

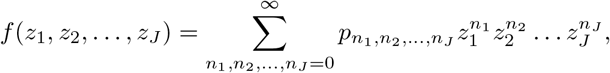

and the Laplace transform of the Poisson kernel *ρ*(*x*_1_, *x*_2_, …, *x*_*J*_) is defined by

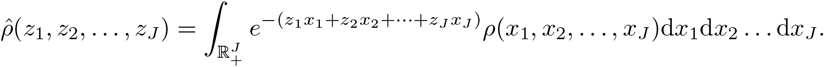

Similarly to the univariate case, in terms of the generating function and Laplace transform, Eq. (40) can be converted into the following equivalent form:

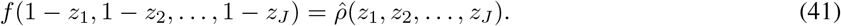

### 5.2 Model of gene regulatory networks

Here we consider the stochastic model of a general gene regulatory network involving protein synthesis, protein degradation, gene state switching, and complex feedback controls. Specifically, we assume that the network involves *J* distinct genes. Each gene can exist in two states: the inactive state *G*_*j*_ and the active state 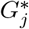. The protein associated with the *j*th gene is denoted by *P*_*j*_, and the corresponding protein number is denoted by *n*_*j*_. Then the network can be described by the following reactions:

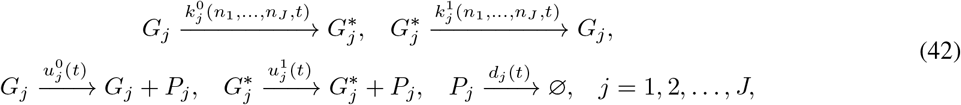

where the first two reactions describe gene state switching, the middle two reactions describe protein synthesis in the two gene states, and the last reaction describes protein degradation. Here we allow all rate parameters to depend on time *t*. Due to complex gene regulation mechanisms, the switching rates 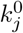 and 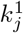 may depend on the numbers of all proteins. For example, if the *i*th gene activates (inhibits) the *j*th gene, then 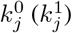 must be an increasing function of *n*_*i*_ since the binding of protein *P*_*i*_ to the promoter of gene *G*_*j*_ activates (inhibits) the expression of this gene.

First we focus on the discrete stochastic model of the gene regulatory network. The microstate of the system can be represented by an ordered 2*J* -tuple (*i*_1_, …, *i*_*J*_, *n*_1_, …, *n*_*J*_), where *i*_*j*_ is the state of the *j*th gene with *i*_*j*_ = 0, 1 corresponding to the inactive and active states, respectively, and *n*_*j*_ is the number of protein *P*_*j*_. Let 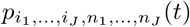 denote the probability of having *n*_*j*_ copies of *P*_*j*_ when the *j*th gene is in state *i*_*j*_, for each *j* = 1, ⋯, *J*. Then the stochastic gene expression dynamics can be modeled discretely by a Markov jump process, and the evolution of the discrete model is governed by the following set of CMEs:

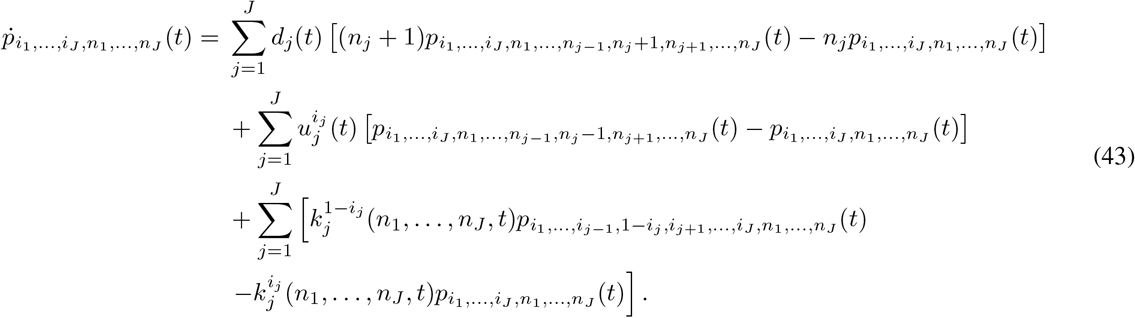

Next we consider the continuous model of the network. In each gene state (*i*_1_, …, *i*_*J*_), the evolution of the network can be modeled by the following set of ODEs:

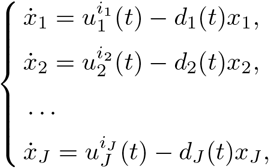

where *x*_*j*_ denotes the abundance of protein *P*_*j*_, and the two terms on the right-hand side describe synthesis and degradation of all proteins involved in the network. Since the genes can switch between different states, the whole gene expression dynamics can be modeled continuously as a switching system of ODEs. Let 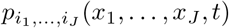 denote the joint probability density of the numbers of all proteins in the gene state (*i*_1_, …, *i*_*J*_). Then the evolution of the continuous model is governed by the Kolmogorov forward equation

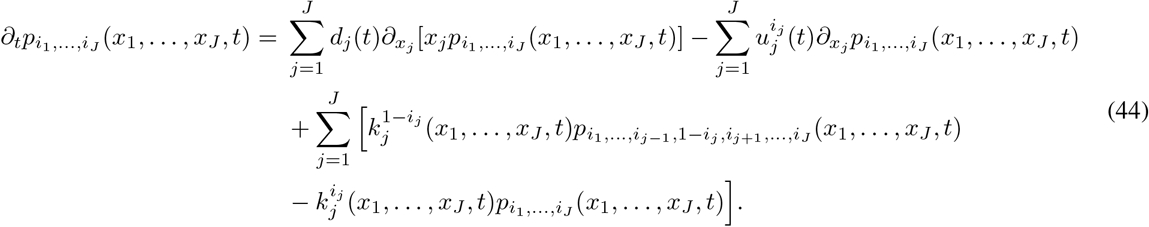

Similarly to autoregulatory systems, we next prove that the joint distributions of the discrete and continuous models are approximately related by the multivariate Poisson representation when the numbers of all proteins are large. To this end, we introduce the system size *K* with *K* → ∞ corresponding to a macroscopic scale. Moreover, we assume that the synthesis rates of all proteins scale with *K* as 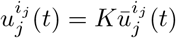, with 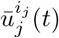 being independent of *K*, and we assume that the switching rates for all genes are functions of the protein concentrations, i.e.

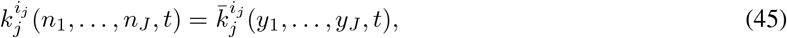

where *y*_*j*_ = *n*_*j*_*/K* is the concentration of *P*_*j*_, and 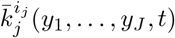 (*y*_1_, …, *y*_*J*_, *t*) are functions independent of *K*.

Next we derive the dynamic equation of protein concentrations from the discrete model. Let 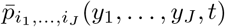 denote the joint probability density of the concentrations of all proteins in the gene state (*i*_1_, …, *i*_*J*_). When *K* ≫ 1, the probability density 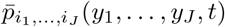 of protein concentrations and the probability distribution 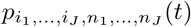 of protein numbers (for the discrete model) should be related by

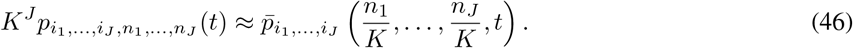

Applying this relation and taking *K* → ∞ in Eq. (43), we find that the evolution of protein concentrations is governed by

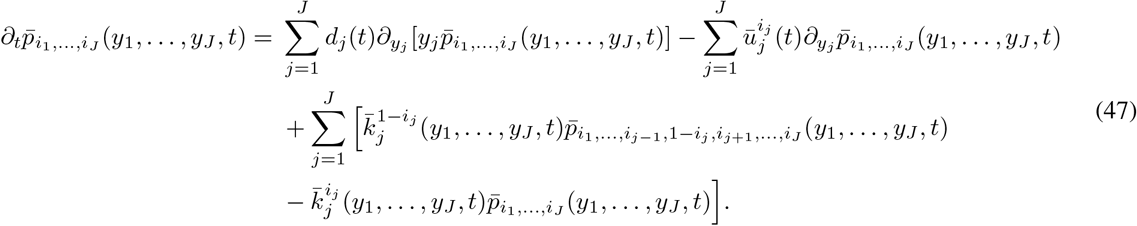

Recall that the dynamics of the continuous model of the network is described by Eq. (44). Comparing Eq. (47) with Eq. (44), it is easy to see that the probability density 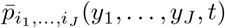 of protein concentrations and the probability density 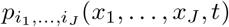 of protein numbers (for the continuous model) are related by

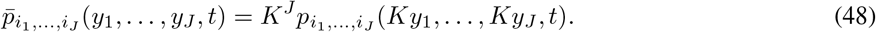

To proceed, let 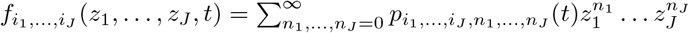 be the generating function of the discrete distribution 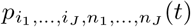. When *K* ≫ 1, applying Eqs. (46) and (48), we obtain

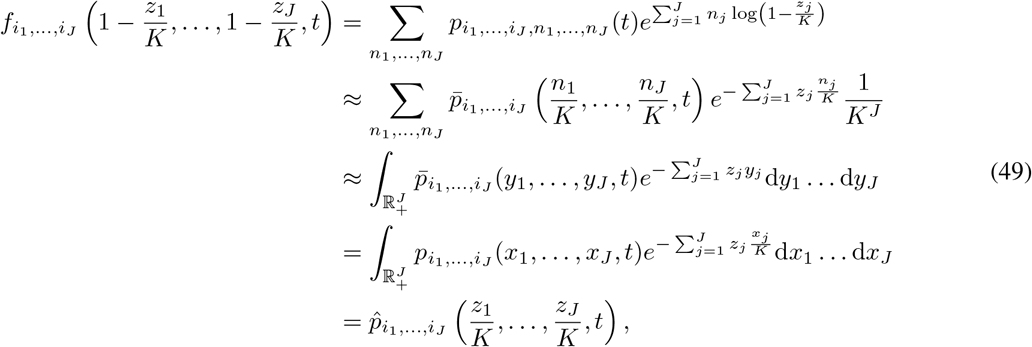

where again we have used the fact that a Riemann sum converges to a Riemann integral as the partition size tends to zero. This clearly shows that when *K* ≫ 1, the generating function of the discrete distribution 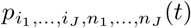 and the Laplace transform of the continuous distribution 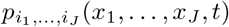 are approximately equal. Hence the solutions of the discrete and continuous models are approximately related by the multivariate Poisson representation in the limit of large protein numbers, i.e.

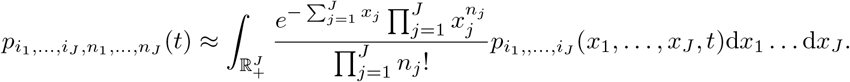

### 5.3 Numerical validation

We then apply our results to a specific gene regulatory network. Here we consider a toggle switch (see Fig. 5(a) for illustration), where two genes repress the expression of each other [83]. The toggle switch can be described by the following reactions:

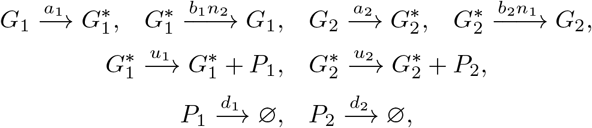

where protein *P*_2_ (*P*_1_) binds to gene *G*_1_ (*G*_2_) and inhibits its expression. The first four reactions describe state switching of the two genes, the next two reactions describe synthesis of the two proteins, and the last two reactions describe degradation of the two proteins.

**Figure 5.**
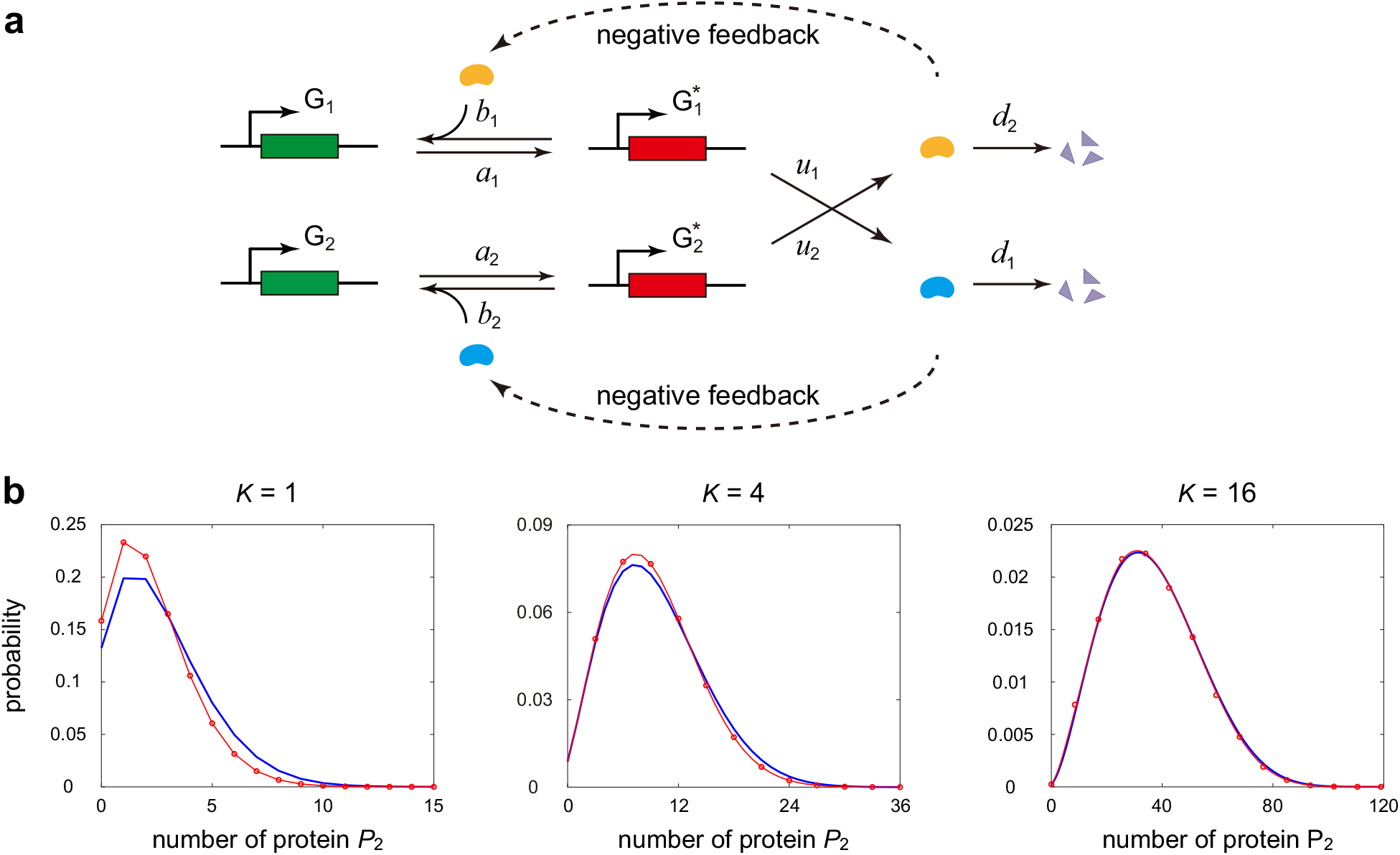
Poisson representation for a toggle switch. **(a)** Schematics of a toggle switch. The protein associated with each gene binds to and inhibits the expression of the other gene. **(b)** Comparison of the steady-state distribution of the discrete model computed using the SSA (blue curves) and the approximate one given by Eq. (50) (red circles and red curves). When computing the approximate distribution, we first calculate the steady-state distribution of the continuous model using the SSA and then construct the approximate distribution using the Poisson representation. The parameters are chosen as *a*_1_ = 2, *b*_1_ = 0.01, *a*_2_ = 3, *b*_2_ = 2.5, *ū*_1_ = 2, *ū*_2_ = 6, *d*_1_ = 1, *d*_2_ = 1.

To validate our results, we assume that the synthesis rates scale with the system size *K* as *u*_*j*_ = *ū*_*j*_*K, j* = 1, 2 and the feedback strengths scale with *K* as 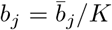, *j* = 1, 2, where we regard *ū*_*i*_, *d*_*i*_,*a*_*i*_, and 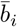 as constants. For simplicity, we focus on the steady-state protein number distributions of the discrete and continuous models, where the continuous model can be described by the following switching system of ODEs:

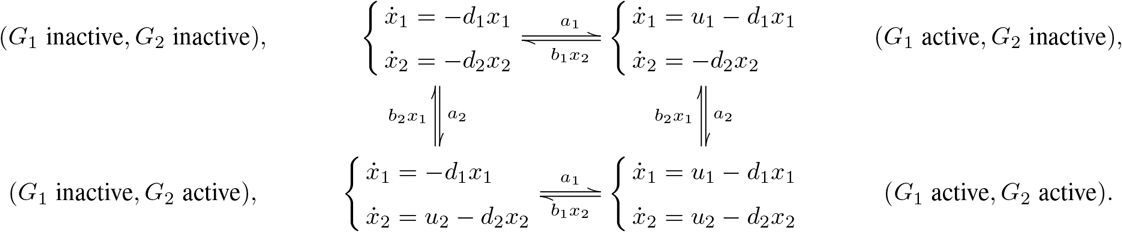

According to our results, the steady-state distributions, 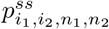 and 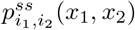, of the discrete and continuous models are approximately related by the two-dimensional Poisson representation when *K* ≫ 1, i.e.

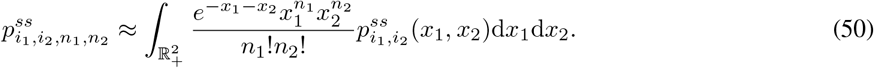

To test this, we compare the steady-state approximate solution of the discrete model generated from the Poisson representation with the numerical one obtained from the SSA. In the Poisson representation, the Poisson kernel is chosen to be the steady-state solution of the continuous model obtained from the SSA. For convenience of comparison, here we only focus on the marginal distribution of protein *P*_2_ (see Fig. 5(b)). It is clear that the Poisson representation fails to connect the discrete and continuous models when protein numbers are small, but it indeed links the two models when protein numbers are large.

## 6 Conclusions and discussion

In this paper, we systematically explore when the Poisson representation can be used to connect the distributions of the discrete and continuous models for stochastic gene regulatory networks. We found that if the gene of interest is unregulated, then the mRNA or protein number distributions for the discrete and continuous models are always related by the Poisson representation, whenever (i) the initial distributions for the two models are related by the Poisson representation and (ii) the burst size distributions for the two models are also related by the Poisson representation. This holds for any unregulated system with complex promoter switching mechanisms, bursty production of the gene product, and even upstream cellular drives. If the distributions for the two models are related by the Poisson representation and the exact distribution for one model is known, then we can use this relation to recover the exact distribution for the other model. This technique was then used to obtain some new analytical results including (i) the steady-state solution for a multi-state bursty model, (ii) the time-dependent solution for a two-state bursty model, and (iii) the time-dependent solution for a coupled dynamic model of gene expression and cell size.

We next examined whether the Poisson representation also plays a crucial role for regulated systems. For an autoregulated gene, we showed by a simple example that the protein number distributions for the discrete and continuous models are not exactly related by the Poisson representation. However, by introducing a macroscopic scaling parameter, we found that the discrete and continuous models are approximately related by the Poisson representation in the limit of large protein numbers. As an application, we derived the approximate distribution for the discrete model of an autoregulatory gene circuit with complex feedback regulation. More interestingly, we generalized these results to a complex regulatory networks with multiple genes. Similarly, we revealed that for an arbitrary gene regulatory network, the joint protein number distributions for the discrete and continuous models are approximately related by the Poisson representation in the limit of large protein numbers.

In summary, the present work shows that the Poisson representation serves as a powerful tool to investigate the dynamic properties of stochastic gene regulatory networks, but it should be used with caution when the molecule numbers of some proteins are very small. We anticipate that the results of the present paper can be extended to non-Markovian [84] and delayed [85] models of gene regulatory networks. It would be also interesting to study whether the concept of Poisson representation and the results of the present paper can be generalized to spatially resolved gene expression models [86].

## Data Availability Statement

The data that supports the findings of this study are available within the article.

## Acknowledgements

The authors thank Dr. Bingjie Wu for valuable suggestions. C. J. acknowledges support from National Natural Science Foundation of China with grant No. U1930402, No. 12271020, and No. 12131005.

## Appendix A The proof of Theorem 1

For the gene expression model described by Eq. (14), the probability distribution *p*_*i,n*_(*t*) of the discrete model satisfies the set of CMEs

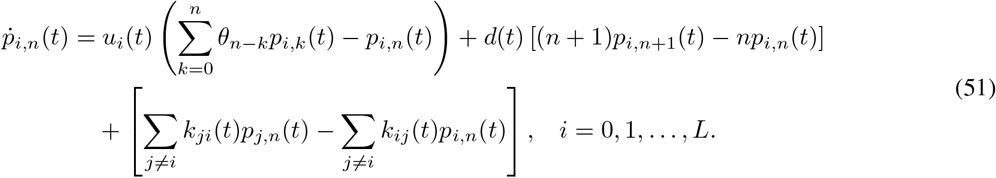

To proceed, let 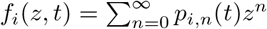 denote the generating function of *p*_*i,n*_(*t*) and let 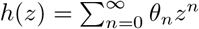 denote the generating function of the burst size distribution *θ*_*n*_ of the discrete model. Note that

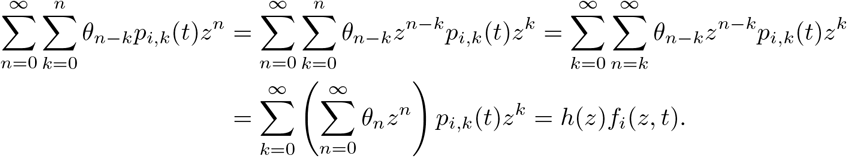

Multiplying *z*^*n*^ on both sides of Eq. (51) and then summing over *n*, we obtain

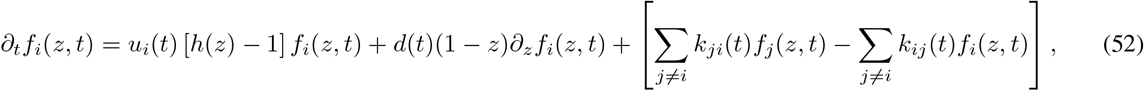

subject to the initial condition

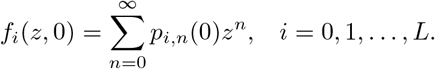

From Eq. (2), the generating function *f*_*i*_(*z, t*) and the Poisson kernel *ρ*_*i*_(*x, t*) of the discrete distribution *p*_*i,n*_(*t*) are related by 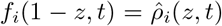, where 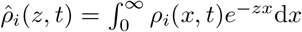 denotes the Laplace transform of *ρ*_*i*_(*x, t*). It then follows from Eq. (52) that the evolution of 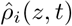 is given by

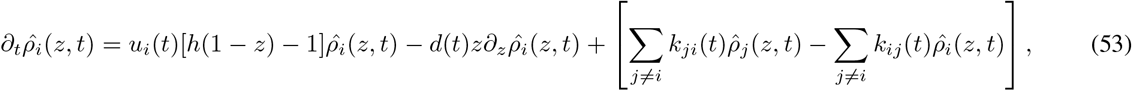

subject to the initial conditions

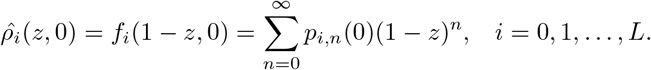

On the other hand, the probability distribution *p*_*i*_(*x, t*) of the continuous model satisfies the Kolmogorov forward equation

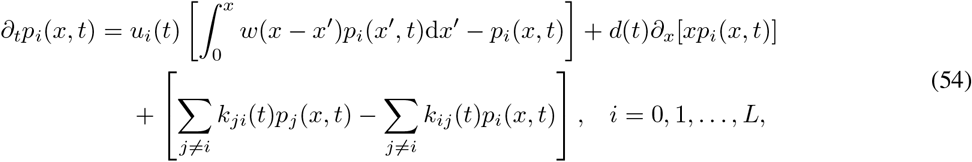

where *w*(*x*) is the burst size distribution of the continuous model. Recall that the burst size distributions of the discrete and continuous models are related by *h*(1 − *z*) = *ŵ* (*z*), where 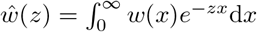 denotes the Laplace transform of *w*(*x*). Taking the Laplace transform with respect to *x* on both sides of Eq. (54), it is easy to check that the evolution of 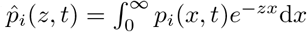 is governed by

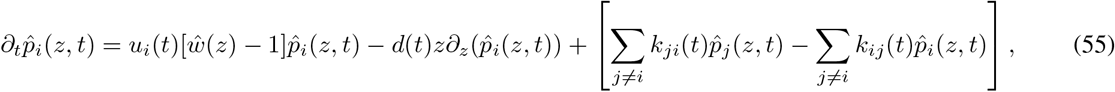

subject to the initial conditions

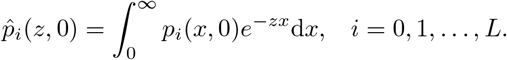

Here we assume that for each gene state *G*_*i*_, the initial distributions of the discrete and continuous models are related by the Poisson representation, i.e.

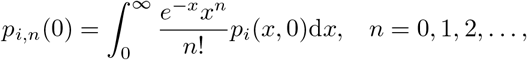

which means that 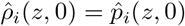. Comparing Eqs. (53) and (55), it is easy to see that 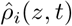 and 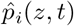 satisfy the same initial value problem. For the initial value problem of a linear system of first-order partial differential equations, the well-known Cauchy-Kovalevskaya theorem [87] states that if the initial conditions of the system of partial differential equations are restricted on a hypersurface and if the coefficients and initial conditions are sufficiently smooth, then there exists a unique solution in the hypersurface. This guarantees the existence and uniqueness of the solution of Eq. (55), which shows that

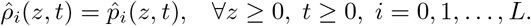

Finally, taking the inverse Laplace transform yields

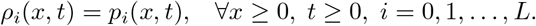

## Appendix B Derivation of Eq. (39)

Recall that for autoregulatory gene circuit whose reactions are given by Eq. (34), the steady-state distribution for the continuous model is given by

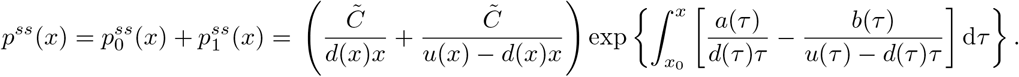

Next we compute the integral in the above equation when

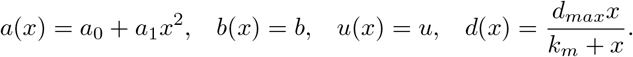

First, it is easy to check that

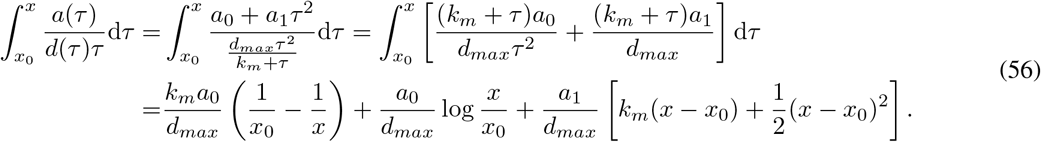

To proceed, note that

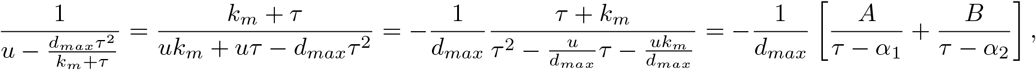

where *α*_1_, *α*_2_, *A*, and *B* are four constants given by

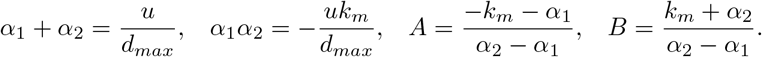

Then we have

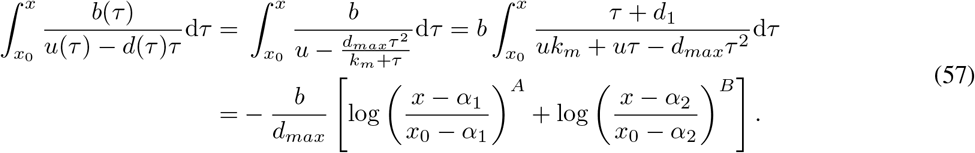

Inserting Eqs. (56) and (57) into Eq. (37) in the main text, we finally obtain Eq. (39).

